# Structural and Functional Comparison of SARS-CoV-2-Spike Receptor Binding Domain Produced in *Pichia pastoris* and Mammalian Cells

**DOI:** 10.1101/2020.09.17.300335

**Authors:** Argentinian AntiCovid Consortium, Claudia R. Arbeitman, Gabriela Auge, Matías Blaustein, Luis Bredeston, Enrique S. Corapi, Patricio O. Craig, Leandro A. Cossio, Liliana Dain, Cecilia D’Alessio, Fernanda Elias, Natalia B. Fernández, Javier Gasulla, Natalia Gorojovsky, Gustavo E. Gudesblat, María G. Herrera, Lorena I. Ibañez, Tommy Idrovo, Matías Iglesias Randon, Laura Kamenetzky, Alejandro D. Nadra, Diego G. Noseda, Carlos H. Paván, María F. Pavan, María F. Pignataro, Ernesto Roman, Lucas A. M. Ruberto, Natalia Rubinstein, Javier Santos, Francisco Velazquez Duarte, Alicia M. Zelada

**Affiliations:** Consejo Nacional de Investigaciones Científicas y Técnicas (CONICET). Godoy Cruz 2290 C1425FQB, Buenos Aires, Argentina; GIBIO-Universidad Tecnológica Nacional-Facultad Regional Buenos Aires. Medrano 951 C1179AAQ, Buenos Aires, Argentina; Theoretical Physics and Center of Interdisciplinary Nanostructure Science and Technology, Universität Kassel. Heinrich-Plett-Str. 40, 34132, Kassel, Germany; Universidad de Buenos Aires. Facultad de Ciencias Exactas y Naturales. Departamento de Fisiología y Biología Molecular y Celular. Instituto de Biociencias, Biotecnología y Biología Traslacional (iB3). Buenos Aires, Argentina; Universidad de Buenos Aires. Facultad de Farmacia y Bioquímica. Departamento de Química Biológica. Junín 965 C1113AAD. Buenos Aires, Argentina; CONICET-Universidad de Buenos Aires. Instituto de Química y Fisicoquímica Biológicas. (IQUIFIB). Buenos Aires, Argentina; Universidad de Buenos Aires. Facultad de Ciencias Exactas y Naturales. Departamento de Química Biológica. Buenos Aires, Argentina; CONICET-Universidad de Buenos Aires. Instituto de Química Biológica de la Facultad de Ciencias Exactas y Naturales (IQUIBICEN). Buenos Aires, Argentina; Universidad de Buenos Aires. Facultad de Ciencias Exactas y Naturales. Departamento de Fisiología y Biología Molecular y Celular. Laboratorio de Agrobiotecnología. Buenos Aires, Argentina; Centro Nacional de Genética Médica, Avda Las Heras 2670, 3er piso, C1425ASP, Buenos Aires, Argentina; Instituto de Ciencia y Tecnología Dr. César Milstein (Consejo Nacional de Investigaciones Científicas y Técnicas-Fundación Pablo Cassará). Saladillo 2468 C1440FFX, Buenos Aires, Argentina; Universidad Nacional de la Plata-CONICET. Centro de Investigaciones del Medio Ambiente (CIM). La Plata, Argentina.; Consejo Nacional de Investigaciones Científicas y Técnicas (CONICET). Instituto de Ciencia y Tecnología Dr. César Milstein. Saladillo 2468, C1440FFX, Buenos Aires, Argentina.; CONICET-Universidad de Buenos Aires. Instituto de Química Física de los Materiales, Medio Ambiente y Energía (INQUIMAE), Buenos Aires, Argentina; CONICET-Universidad de Buenos Aires. Facultad de Medicina. Instituto de Investigaciones en Microbiología y Parasitología (IMPaM). Buenos Aires, Argentina; Universidad Nacional de San Martín-CONICET. Instituto de Investigaciones Biotecnológicas (IIBio). San Martín, Buenos Aires, Argentina; Universidad de Buenos Aires. Facultad de Farmacia y Bioquímica. Buenos Aires, Argentina.; CONICET-Universidad de Buenos Aires. Facultad de Medicina. LANAIS-PROEM. Instituto de Química y Fisicoquímica Biológicas. (IQUIFIB). Buenos Aires, Argentina; Universidad de Buenos Aires. Facultad de Farmacia y Bioquímica. Departamento de Microbiología, Inmunología, Biotecnología y Genética. Buenos Aires, Argentina; CONICET-Universidad de Buenos Aires. Facultad de Farmacia y Bioquímica. Instituto de Nanobiotecnología (NANOBIOTEC). Buenos Aires. Argentina; Instituto Antártico Argentino. Ministerio de Relaciones Exteriores y Culto. Buenos Aires, Argentina; CONICET-Universidad de Buenos Aires. Instituto de Biodiversidad y Biología Experimental y Aplicada (IBBEA). Buenos Aires, Argentina

## Abstract

The yeast *Pichia pastoris* is a cost-effective and easily scalable system for recombinant protein production. In this work we compared the conformation of the receptor binding domain (RBD) from SARS-CoV-2 Spike protein expressed in *P. pastoris* and in the well established HEK-293T mammalian cell system. RBD obtained from both yeast and mammalian cells was properly folded, as indicated by UV-absorption, circular dichroism and tryptophan fluorescence. They also had similar stability, as indicated by temperature-induced unfolding (observed *T*_m_ were 50 °C and 52 °C for RBD produced in *P. pastoris* and HEK-293T cells, respectively). Moreover, the stability of both variants was similarly reduced when the ionic strength was increased, in agreement with a computational analysis predicting that a set of ionic interactions may stabilize RBD structure. Further characterization by HPLC, size-exclusion chromatography and mass spectrometry revealed a higher heterogeneity of RBD expressed in *P. pastoris* relative to that produced in HEK-293T cells, which disappeared after enzymatic removal of glycans. The production of RBD in *P. pastoris* was scaled-up in a bioreactor, with yields above 45 mg/L of 90% pure protein, thus potentially allowing large scale immunizations to produce neutralizing antibodies, as well as the large scale production of serological tests for SARS-CoV-2.

## Introduction

The COVID-19 outbreak was first recognized in December 2019 in Wuhan, China^1^. Since then, this virus has spread to all parts of the world, resulting in a total of 29415168 infected individuals and 931934 deaths by September 14^th^, 2020 (https://www.coronatracker.com). The causative agent is a coronavirus that causes a severe acute respiratory syndrome (SARS). This SARS-related coronavirus (SARSr-CoV) has been designated as SARS-CoV-2.

Coronaviruses are enveloped non-segmented positive sense RNA viruses^2^ that have four open reading frames (ORFs) for structural proteins -Spike, Envelope, Membrane, and Nucleocapsid-^3, 4^, from which Spike is the primary determinant of CoVs tropism. Spike mediates the viral and cellular membrane fusion by binding mainly to the angiotensin-converting enzyme 2 (ACE2), a homologue of ACE^5, 6^.

The SARS-CoV-2 genome has 29903 nucleotides in length^7^, sharing 79% and 50% sequence identity with SARS-CoV-1 and MERS-CoV genomes, respectively^8^. Genetic studies suggest that both viruses originated from bat CoVs^8, 9^, with civet cats as intermediate hosts in the case of SARS-CoV-1^10^, and pangolins in the case of SARS-CoV-2^11, 12^. Among the structural proteins, the Envelope protein has the highest sequence similarity between SARS-CoV-2 and SARS-CoV-1 (96% identity), while the Spike protein, responsible for the interaction with the host receptor, the largest sequence divergence (76% identity with SARS-CoV-1)^13^. It has been suggested that the divergence of Spike could be related to an increased immune pressure^1^. Consistently with the proposed role of pangolins as SARS-CoV-2 intermediate hosts, CoVs from pangolins share the highest genetic similarity with this virus in the region encoding the receptor binding domain (RBD) of the Spike protein^11, 14^.

Due to its important role for SARS-CoV-2 entry into the host cell, Spike is the most studied protein of this virus. This transmembrane glycosylated protein is composed of 1273 amino acid assemblies as a homotrimer that forms spikes that protrude from the virus envelope. Spike has two domains, named S1 and S2. Residues 319-591 from S1 correspond to the RBD, responsible for the interaction with ACE2^15^. RBD binds with high affinity to the ACE2, located on the outer surface of the cell membrane, which acts as a SARS-CoV-2 receptor since it mediates the fusion of the virus to the cell membrane^16^. Spike also includes a transmembrane domain and a fusion peptide^17^.

Molecular dynamics simulations suggest that internal motions of the Spike trimer are important to expose the RBD domain so it can interact with the target receptor. However, part of the time RBD domain is hidden within the rest of the Spike protein, and this process is mediated by protein motions of high amplitude. The structure of RBD-ACE2 protein complex and the structure of Spike (as the full-length and trimeric form of the protein) were determined by X-ray crystallography and cryo-EM^16, 18–20^.

RBD is a protein domain of 220 residues, it has nine cysteine residues (eight of them forming disulfide bonds) (**Figure 1**) and two *N-*glycosylation sites (N331 and N343). The addition of glycan moieties might have a relevant role on the *in vivo* protein folding process, on the dynamics, stability and solvent accessibility of RBD and also on its immunogenicity^21, 22^. RBD is not a globular protein domain; it has a central twisted antiparallel beta-sheet formed by five strands decorated with secondary structure elements (short helices and strands) and loops^19^. The secondary structure analysis of the protein shows 12.4% helix, 33.0% sheet, 19.1% turn, and 35.6% coil.

**Figure 1.**
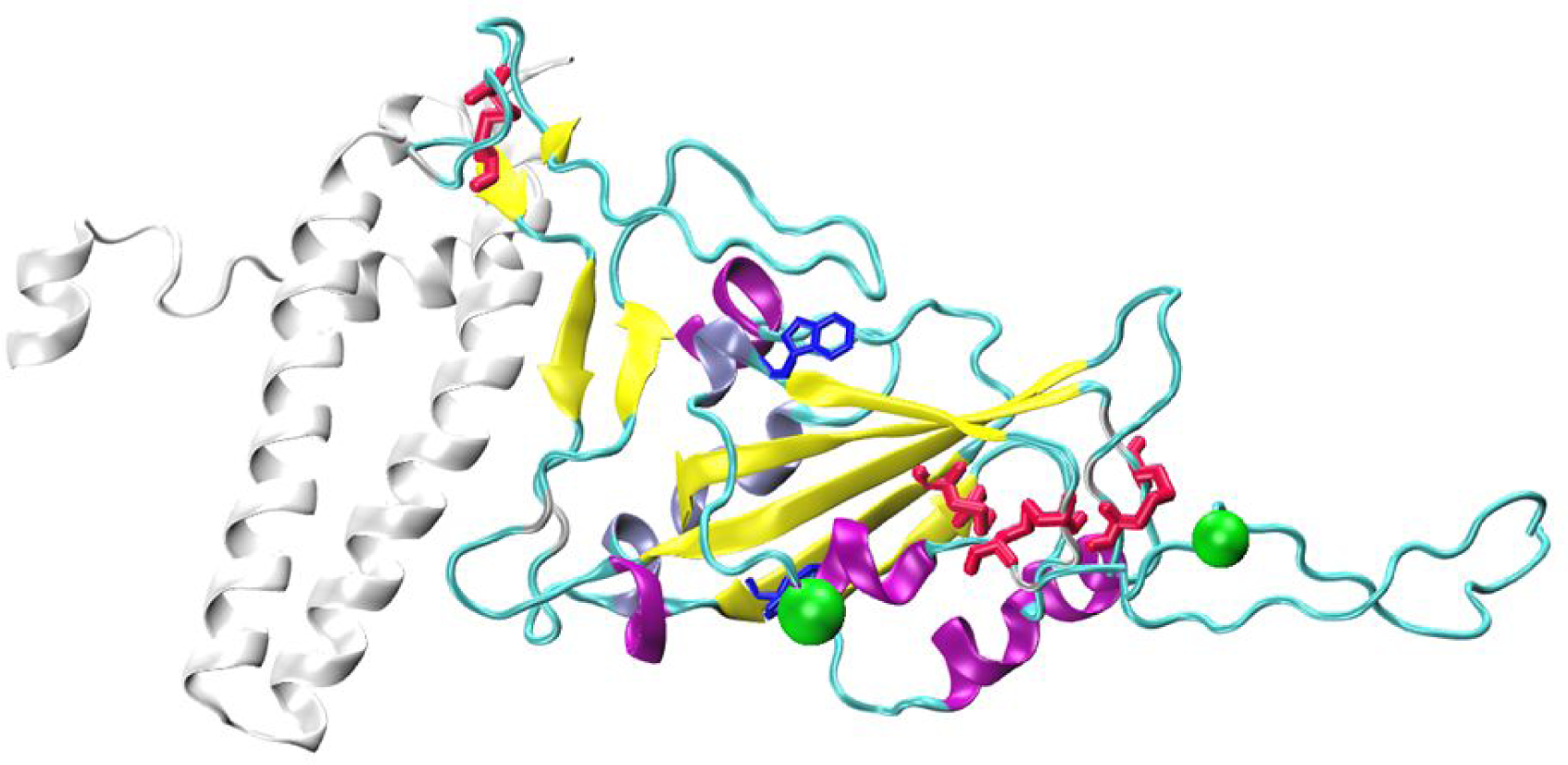
Structure of Sars-Cov-2 Receptor Binding Domain bound to ACE2. The secondary structure elements of RBD are differentially colored (Alpha helices: purple, 3_10 helices: iceblue, beta strands: yellow, and turns/coil: cyan). Disulfide bridges (red) and tryptophan residues (blue) are shown as sticks, while N-glycosylation asparagine residues (green) are shown as VDW spheres. The region of ACE2 encompassing residues 1-115 (colored white) which interacts with RBD is also shown. The structure was generated using PDB structures 6xm0 and 6m0j.

Despite its medium size of 25 kDa, RBD is an example of a challenging protein domain to express in heterologous systems due to its complex topology (Figure 1). Nevertheless, it is of high importance to produce and purify RBD at low-cost and efficiently, since this domain is extensively used for the development of serological test kits as well as an immunogen, both for the production of animal immune sera and for vaccine development^23^. While *E. coli* is a cost-efficient system for the expression of many proteins, it is unlikely to be the case for RBD due to its requirement of disulfide bond formation and glycosylation for its proper expression and folding. For this reason, RBD is usually expressed in mammalian as well as insect cells^18, 24^.

The methylotrophic yeast *Pichia pastoris* is an alternative cost-effective eukaryotic system that allows relatively easy scaling-up of recombinant protein production, and which has previously been used for the expression of SARS-CoV-1 RBD to produce a vaccine^25^. This yeast can use methanol as an exclusive carbon source. This molecule is also an inductor of the strong and tightly regulated AOX1 promoter^26^, which can therefore be used to drive recombinant protein expression. When cultured in bioreactors, *P. pastoris* can reach high cell densities, and more importantly, this organism allows the efficient secretion of recombinant proteins to the culture medium, which contains relatively low levels of endogenous proteins, thus allowing the straightforward purification of recombinant secretory proteins^26^.

In this work we expressed and purified SARS-CoV-2 Spike RBD from two different systems -the yeast *P. pastoris* and mammalian cells- and compared their structure, stability, glycosylation status, and immunogenicity in mice. Our work provides useful insights on the production of a key protein used in diagnosis and therapeutics to fight COVID-19 pandemia.

## Results

### SARS-CoV-2 RBD Protein Sequence analysis

Prior to designing the constructs to express Spike RBD domain from SARS-CoV-2 we looked for possible variation in its coding sequence in genomes publicly available at the Global Initiative for Sharing All Influenza Data (GISAID) database (https://www.gisaid.org)27. From a total of 75355 SARS-CoV-2 genome sequences available at GISAID, 85.8% (64,707 genomes) have 100% coverage of RBD (non truncated Spike proteins) with 100% of amino acid identity to the first published RBD sequence (Uniprot: QHN73795.1)^28^. This data set includes 38 Argentinean SARS-CoV-2 genomes. RBD sequences from the remaining genomes (14.2%) were distributed as follows: 3.5 % (6199/64707) have more than 99% of amino acid sequence identity (up to 2 amino acid substitutions or InDels), 1.4% (925/64707) have more than 80% (up to 44 amino acid substitutions or InDels) and only 300 genomes have a lower amino acid identity relative to the first published sequence. Thus, we considered appropriate to express the predominant RBD form, spanning from residue 319 to 537 of Spike protein, which consists of a relatively compact domain, and includes a slightly disordered C-terminal stretch useful for protein engineering (**Figure 1**).

### Expression of RBD in mammalian and yeast cells

The expression of RBD in mammalian cells (HEK-293T cell line) and in *P. pastoris* yielded significant quantities of protein (∼5 and 10-13 mg L^-1^ of cell culture, respectively, at a laboratory scale). In both cases, the recombinant protein was fused to appropriate secretion signal peptides, IL2 export signal peptide for HEK-293T expression and *Saccharomyces cerevisiae* α-factor secretion signal for *P. pastoris* expression. Both secretion signals allowed the recovery of mature RBD from cell culture supernatants.

Since RBD expressed in both eukaryotic systems included a C-terminal His tag, similar purification protocols were used in both cases. However, given that the physico-chemical conditions required for optimal growth of mammalian and yeast cells were completely different (HEK-293T cells were grown at 37 °C in a medium buffered to pH 7.4, while *P. pastoris* were grown at 28 °C, buffered to pH 6.0), the covalent structure, intactness, conformation, post translational modifications and stability of RBD might still differ depending on the expression system used. Additionally, both strategies involved the accumulation of soluble RBD in the supernatant, which can pose an extra challenge for unstable proteins. For these reasons, it was crucial to evaluate parameters such as protein aggregation, oxidation, and possible alterations in disulfide bond patterns of proteins obtained from the different media.

NTA-Ni^2+^-purified RBD from both HEK-293T and yeast exhibited high purity (>90%), as judged by SDS-PAGE analysis (**Figure 2A**). RBD from HEK-293T cells migrated as a 35 KDa single-smear band in SDS-PAGE 12%, while RBD produced in *P. pastoris* migrated as one highly diffuse and more abundant band of ∼45-40 kDa, and a less abundant band of ∼35 kDa, the latter similar to that of RBD produced in HEK-293T cells. HPLC profile analysis revealed that RBD produced in HEK-293T cells is highly homogeneous, as shown by its elution as a sharp peak at 48-49 % of acetonitrile in a reverse phase C18 column, while RBD produced in *P. pastoris* showed a considerably broader peak, although it eluted at very similar acetonitrile concentration. In addition, two very small peaks appeared in the chromatogram of RBD produced in *P. pastoris*.

**Figure 2.**
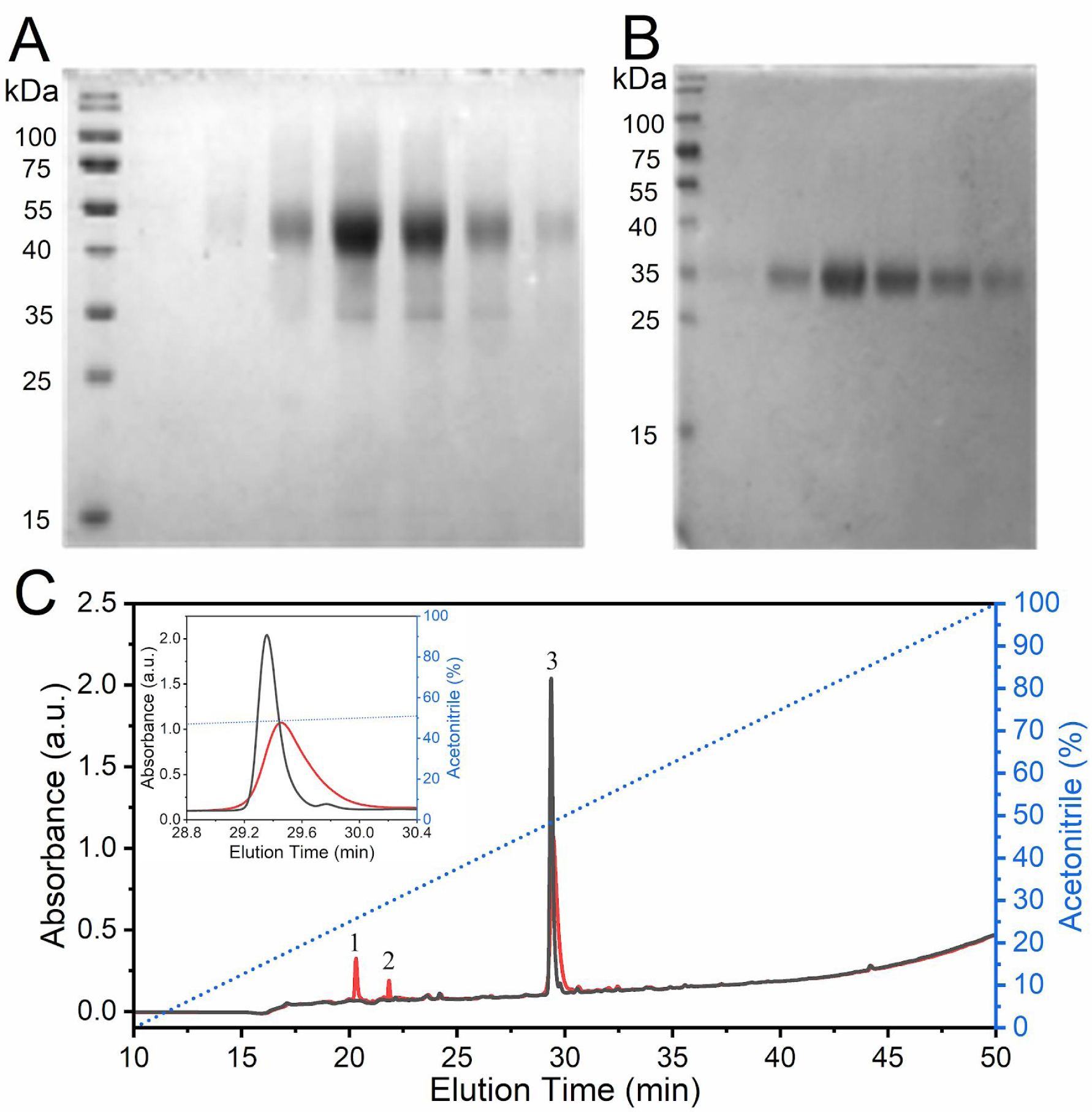
Analysis by SDS-PAGE and RP-HPLC of RBD produced in *P. pastoris* or HEK-293T, and purified by NTA-Ni^2+^. Analysis of recombinant RBD fractions eluted from a NTA-Ni^2+^ column by 300 mM imidazole after purification from supernatants of a *P. pastoris* culture (A) or of HEK-293T cells (B). (C) Reverse Phase HPLC Analysis of RBD. Profiles for RBD produced in *P. pastoris* (red) and HEK-293T mammalian cells (black). The inset shows the expanded region of the chromatogram where the highest peaks eluted. The dashed blue line indicates the variation of acetonitrile (% v/v) during the experiments. Peaks 1, 2 and 3 from RBD produced in *P. pastoris* correspond to areas of 10.1, 2.8 and 87.1%, respectively.

The area corresponding to the full-length protein was approximately 87%.

The SDS-PAGE analysis of RBD purified from yeast and mammalian cell culture supernatants suggested the existence of glycosylation as the main post-traslational modification in RBD, as its theoretical mass (deduced from the amino acid sequence) is ∼26 kDa (**Figure 2**), while both recombinant RBD forms migrated as products of more than 32-35 kDa. This was expected, since two *N*-glycosylation consensus sequences (NIT and NAT) are present at RBD N-terminal region. RBD from SARS-CoV-1 also bears three glycosylation sites at its N-terminal region, and was found to be glycosylated^25^.

Even though Coomassie Blue staining showed heterogeneity in protein size, incubation with PNGaseF, a peptide-endoglycanase that removes high mannose, complex and hybrid N-glycans from proteins, homogenized all isoforms to a sharper band of ∼25-26 kDa, compatible with the predicted MW of deglycosylated RBD (26.5 kDa, **Figure 3**). The decrease in the molecular mass of RBD by endoglycanase digestion confirmed the existence of *N*-glycosylations in both proteins. Moreover, glycans from. *P. pastoris-*RBD - and not from HEK-293T-RBD - were also removed by EndoH, an endoglycanase that eliminates only high-mannose type glycans, which are the expected type in *P. pastoris* yeasts. These results strongly suggest that RBD from mammalian cells bears only complex or hybrid glycans, while RBD from *P. pastoris* only bears high mannose glycans. Moreover, the persistence of two bands in RBD from HEK-293T cells after exhaustive deglycosylation with PNGaseF suggests the existence of heterogeneous *O*-glycosylation, although heterogeneity in amino acid sequence cannot be discarded either.

**Figure 3.**
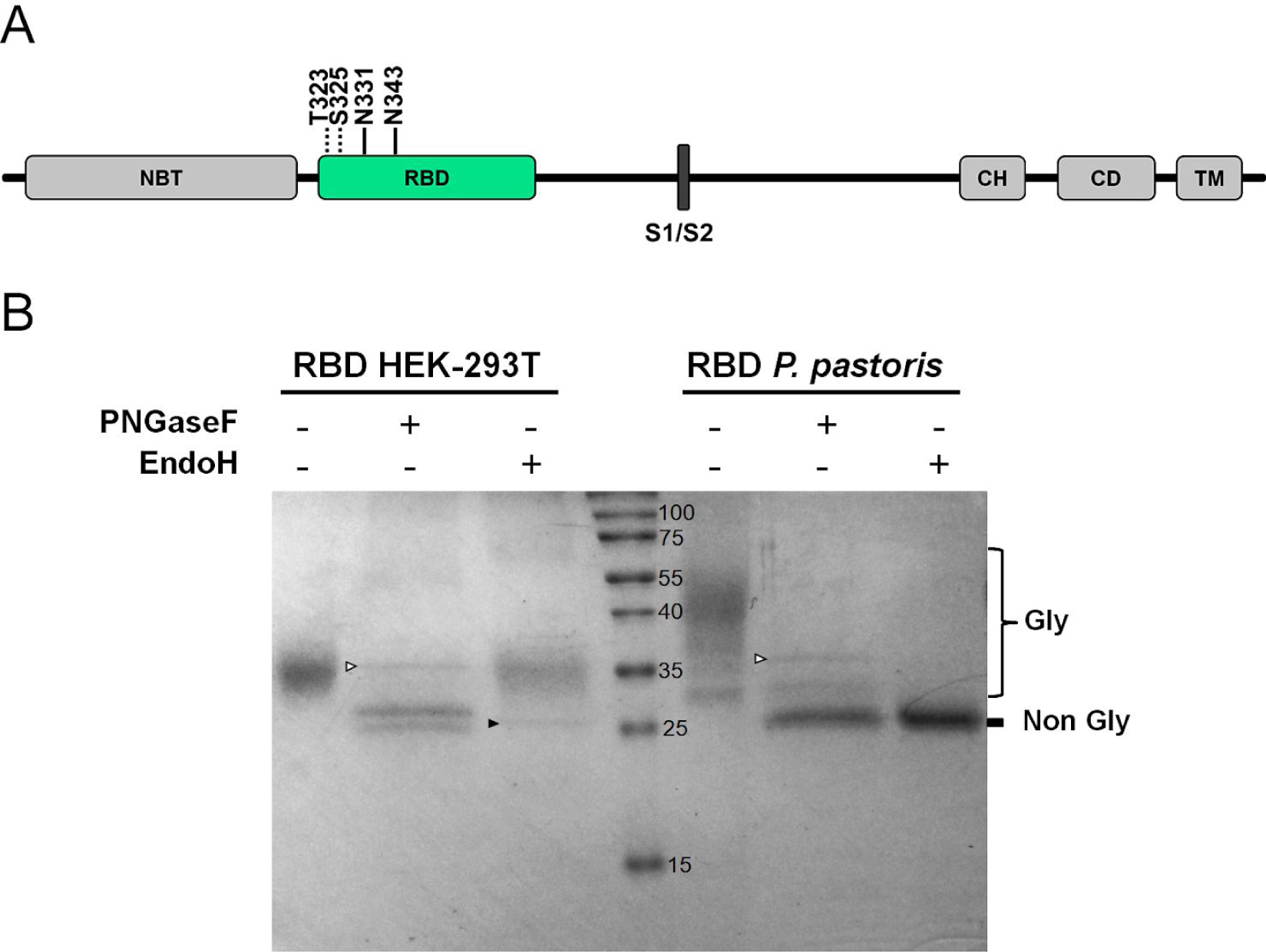
Analysis of the glycosylation status of RBD produced in HEK-293T and *P. pastoris*. (A) Schematic representation of SARS-CoV-2 S glycoprotein. N-terminal domain (NTD), receptor-binding domain (RBD), furin cleavage site (S1/S2), central helix (CH), connector domain (CD), and transmembrane domain (TM) are displayed. Residues involved in RBD glycosylation are shown (*O*- and *N*-glycosylations are indicated by dotted and solid lines, respectively). (B) Endoglycanase treatment of RBD. Purified RBD (3 ug) from mammalian or yeast culture supernatants was denatured 10 min at 100 °C and digested with PNGase F (500 mU) or EndoH (5 mU) during 2 h at 37 °C. Proteins were separated in a 14% SDS-PAGE gel. The positions of non-glycosylated and glycosylated RBD isoforms are indicated. The bands corresponding to PNGaseF (36 KDa) and EndoH (29 KDa) are indicated by empty or full arrowheads, respectively.

The identity of the RBD forms was corroborated by fragmentation, controlled proteolysis, peptide assignment and MS/MS sequencing (MALDI TOF TOF for tryptic peptides analysis). **Figure 4** shows molecular masses and spectra from the intact mass analysis, which agree with values expected for the samples.

**Figure 4.**
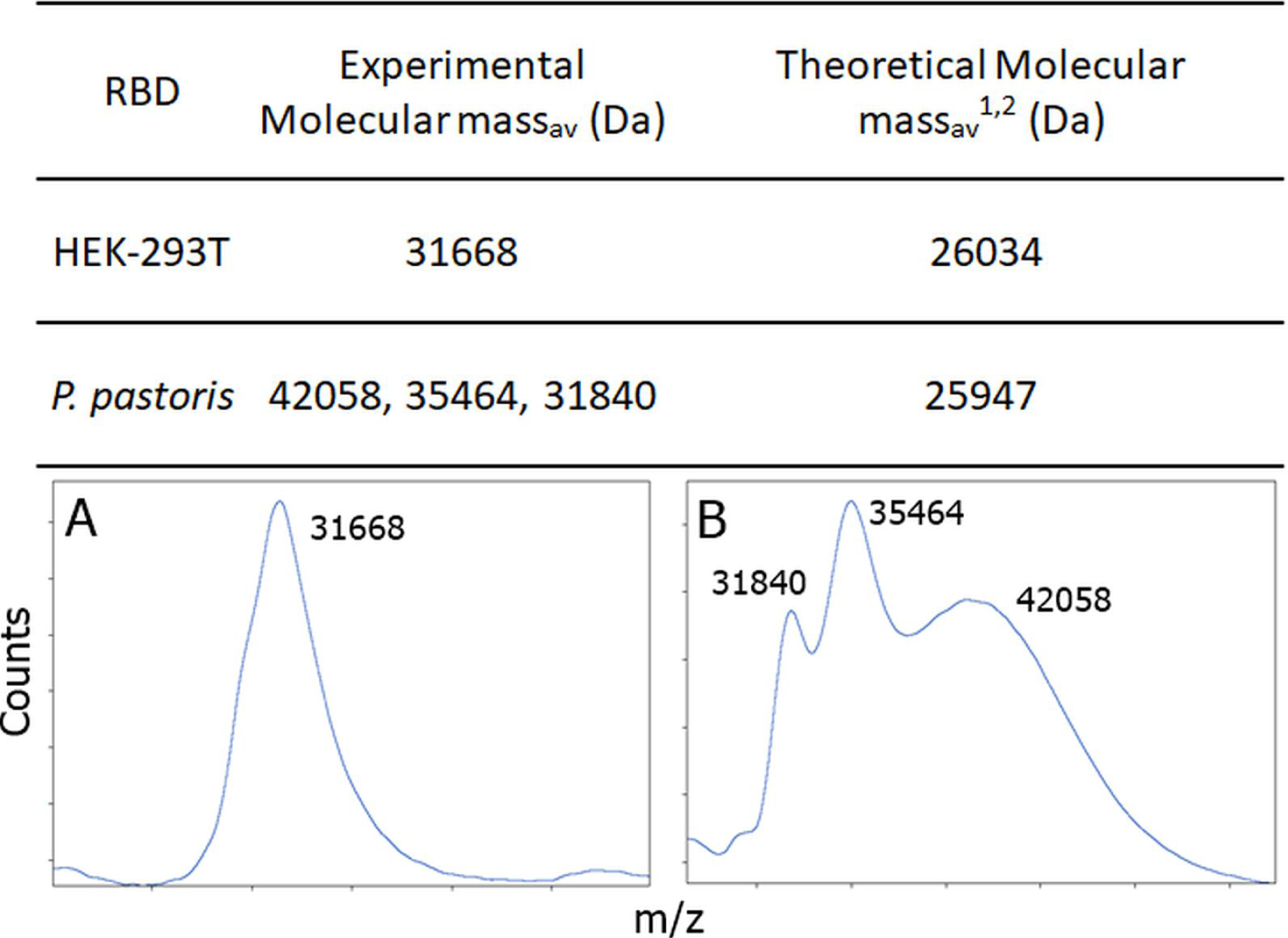
MALDI TOF Spectra for RBD Samples. (A) RBD prepared in HEK-293T and (B) RBD prepared in *P. pastoris*. As expected, the glycosylated species from *P. pastor*is have a broad mass.

**Table 2.**
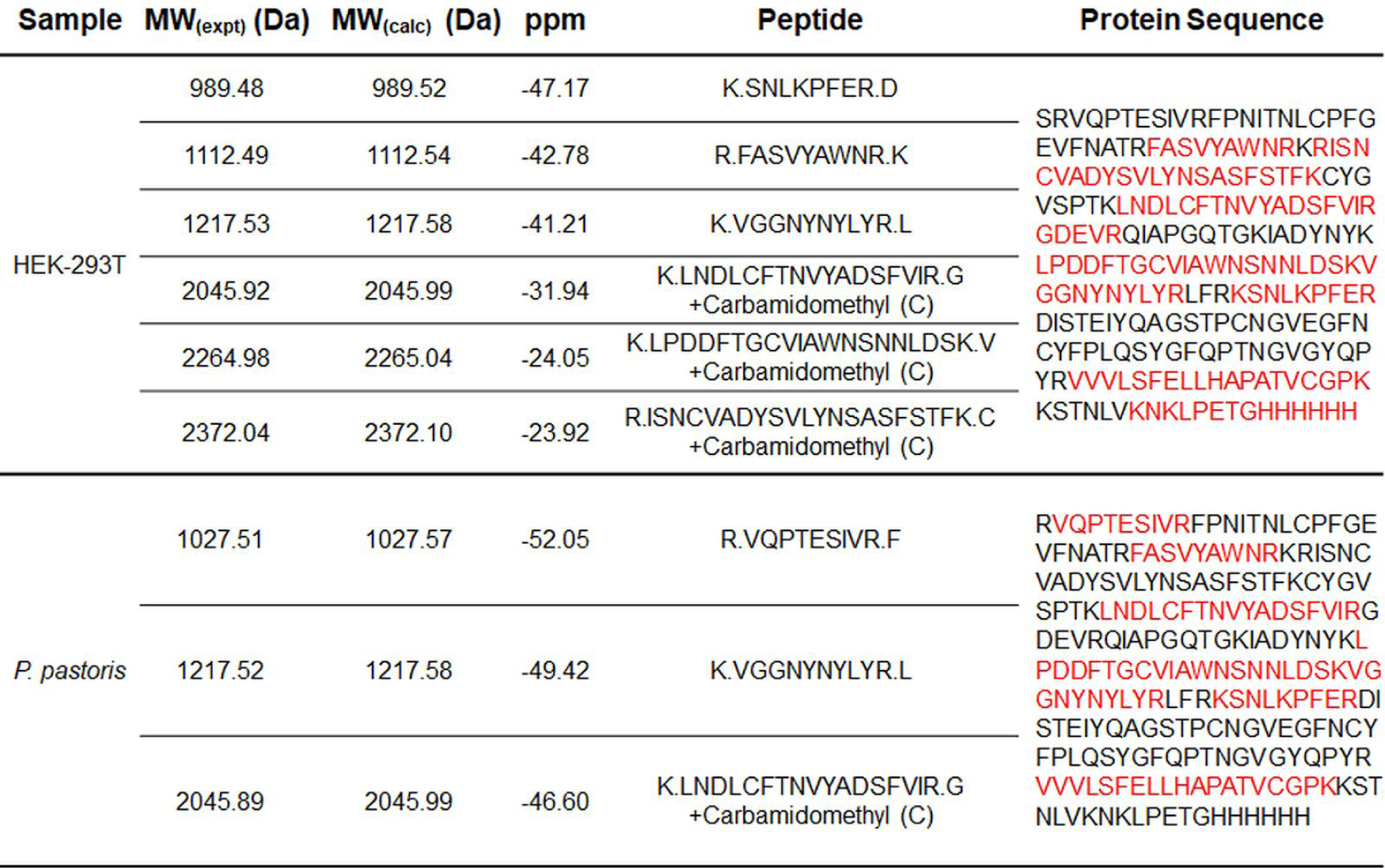
MS/MS Analysis Results for In-solution Digestion of RBD obtained from HEK-293T and from *P. pastoris*. Protein coverage was of ∼40 and ∼60%, for RBD produced in *P. pastoris and* HEK-293T cells, respectively. The experimentally obtained (MW (expt)) and the calculated (MW (calc)) molecular weights of the peptides, and their difference (ppM) are shown. The protein sequence of each construct is shown, red letters indicate the peptides identified either by MS/MS analysis or peptide mass fingerprint (less than 60 ppm error).

The peptide spectrum matches – PSM – with significant score from the MS/MS analysis clearly showed that protein species present in the samples belonged to the RBD from SARS-CoV-2, (**Table 2**). The masses of peptides identified by MS/MS or peptide mass fingerprint (**Table 2**) were in good agreement with those expected from a proteotypic peptide prediction software -PeptideRank^29^, and from the information available from the Peptide Atlas database^^30^^ for peptides identified from SARS-CoV-2. To further validate the PSM findings, the data was also analyzed with COMET at Transproteomic Pipeline (a different MS/MS search engine), which produced similar results^31, 32^.

As expected due to the dispersion in sizes, the FPNITNLCPFGEVFNATR peptide that harbors two glycosylation consensus sequences –NIT and NAT motifs– was not observed in the RBD samples from *P. pastoris* or HEK-293T cells. On the other hand, a deglycosylation of RBD produced in *P. pastoris*, followed by MS/MS analysis revealed a m/z signal only present in that sample, in which two HexNAc moieties identified at the two N residues in FPNITNLCPFGEVFNATR (**Figure S1**).

### Conformational characterization of RBD forms by UV absorption

Different analytical techniques were used to characterize proteins produced in yeast or mammalian cells. The UV absorption spectra of both recombinant proteins were very similar; they are dominated by a high content of tyrosine residues (16 Tyr, 2 Trp, 15 Phe, 4 disulfide bonds), as indicated by bands at approximately 276.0 and 281.0 nm. The absence of light scattering -suggested by the absence of a typical slope between 340 and 300 nm- strongly indicated that the proteins do not form soluble aggregates (**Figure 5**). However, freezing and thawing resulted in protein precipitation when RBD concentration was higher than 80 μM (data not shown).

**Figure 5.**
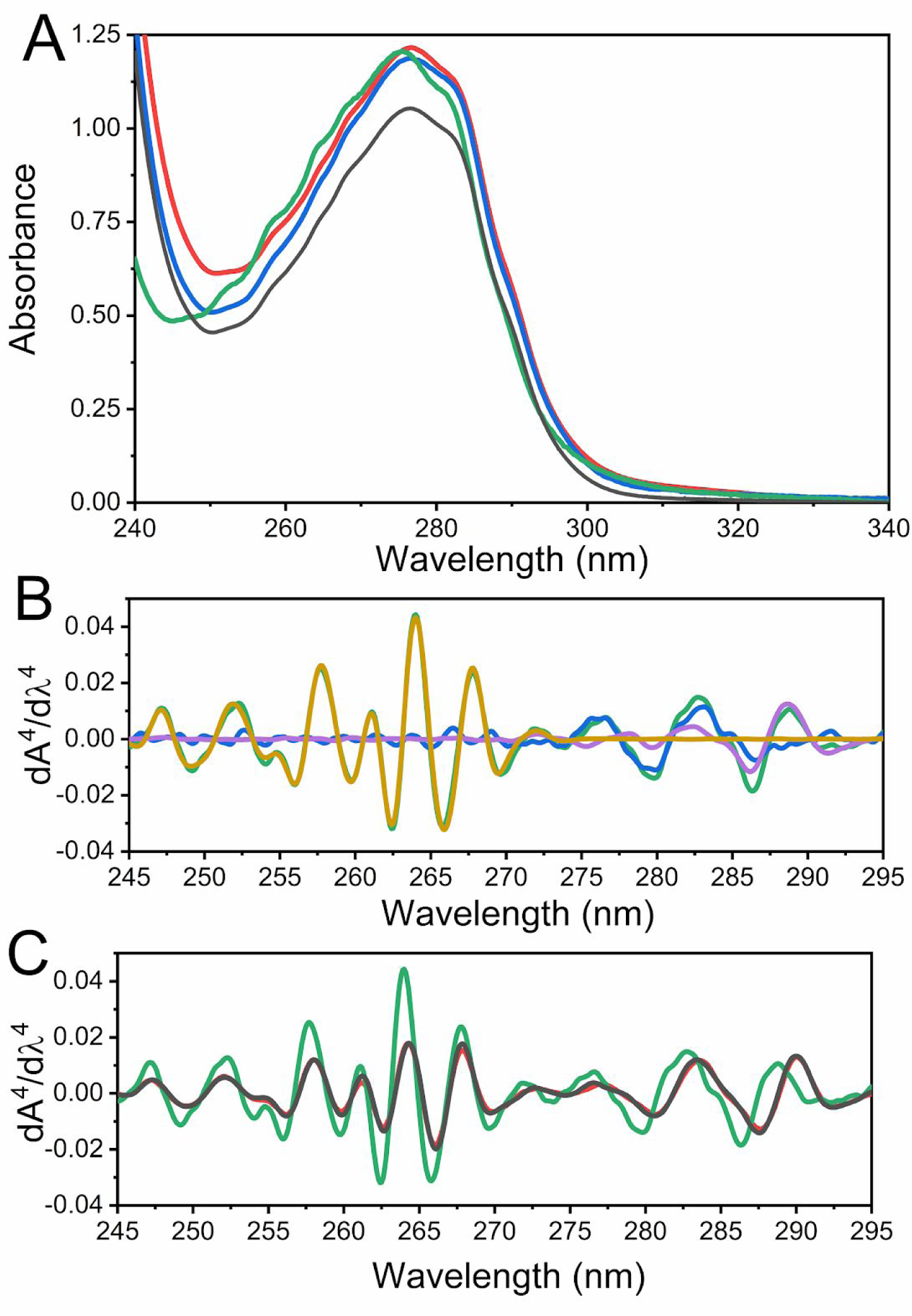
Absorption Spectroscopy. (A) Spectra corresponding to purified RBD produced in HEK-293T cells (black), *P. pastoris* (clones 1 (blue) and 7 (red), and a simulated RBD spectrum (green) (27.5 μM) calculated from its composition of aromatic amino acid (15 Phe, 15 Tyr and 2 Trp in RBD). (B) Fourth derivative spectra corresponding to the aromatic amino acids (Tyr (blue), Trp (violet), Phe (yellow)) and a simulated RBD spectrum (green). (C) Comparison between the fourth derivative spectra from RBD obtained in HEK-293T (black), *P. pastoris* RBD Clone 7 (red) and the simulated spectrum presented in A (green).

The fourth derivative of absorption spectra can be used to evaluate RBD native conformation. Spectra corresponding to RBD produced in HEK-293T cells and *P. pastoris* were superimposable (**Figures 6B and C**), suggesting a similar packing of the aromatic residues. In particular, the positive band at 290.4 nm corresponding to Trp residues observed in the native state of RBD (**Figure 6C**) showed a significant red shift compared to the 288 nm band of N-acetyl-L-tryptophanamide (NATA), suggesting that Trp residues in RBD are not fully exposed to the solvent (**Figure 6B**). Also the negative band at 287.8 nm (a contribution of Tyr and Trp residues) showed a significant red shift compared to that observed for the fully exposed NATA and N-acetyl-L-tyrosinamide, NAYA. Similarly, a band corresponding to Tyr (280.4 nm) showed a shift to 279 nm (NAYA).

**Figure 6.**
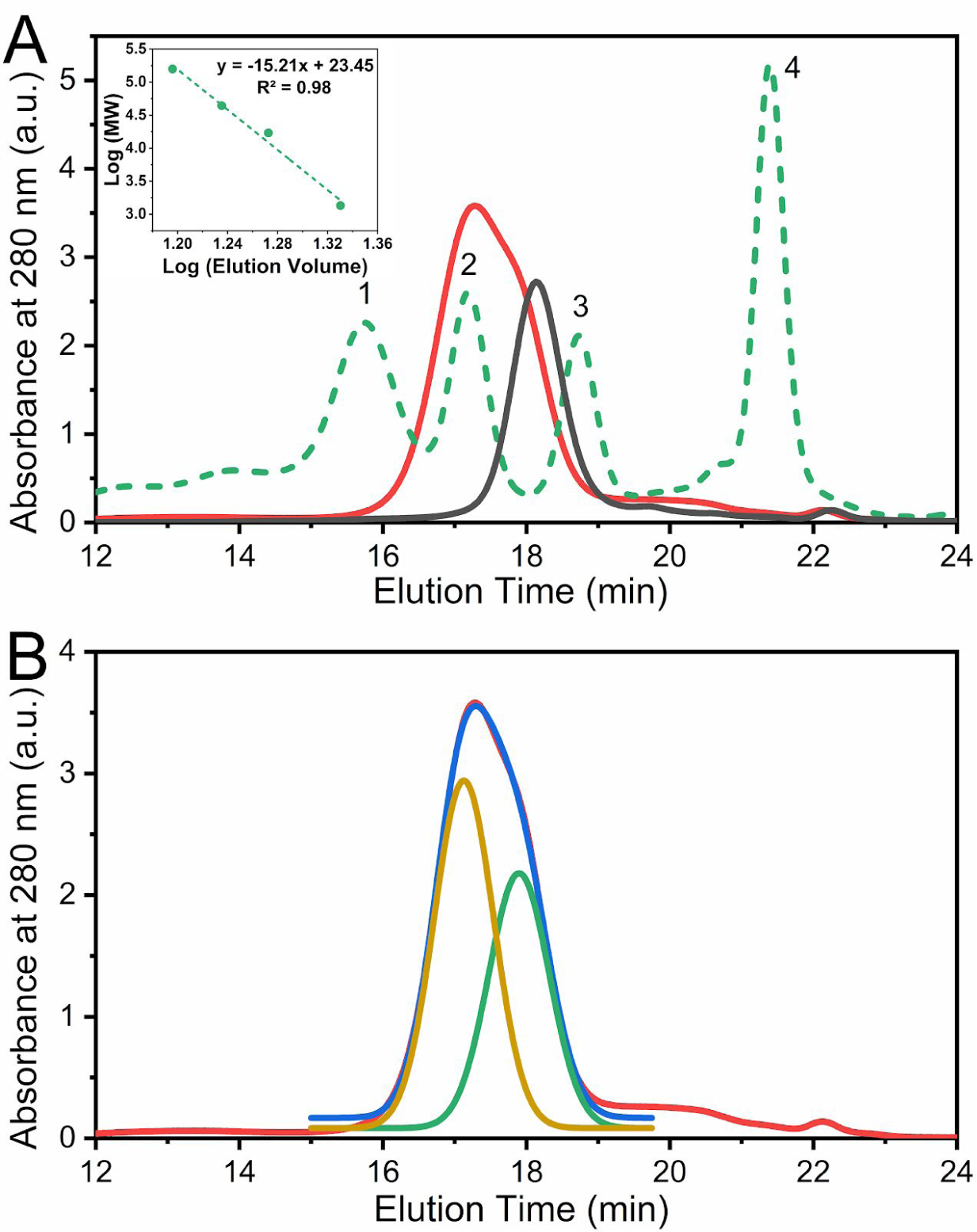
Hydrodynamic Behavior of RBD. (A) SEC-HPLC of RBD produced in *P. pastoris* (red), HEK-293T (black) and molecular weight markers (dashed green line). This analysis was carried out by injecting 50 μL protein aliquots (0.70 and 0.75 mg/mL for RBD produced in *P. pastoris* and HEK-293T, respectively) in 20 mM Tris-HCl, 100 mM NaCl, pH 7.0 buffer. The inset shows the correlation between molecular weight and elution volume obtained from the molecular weight markers: (1) gammaglobulin (158 kDa), (2) ovoalbumin (44 kDa), (3) myoglobin (17 kDa), and (4) vitamin B12 (1350 Da). (B) Deconvolution analysis of the chromatographic profile from RBD from *P. pastoris*. The experimental profile (red), deconvolution of the peak in two different gaussian curves (green and yellow) and the sum of the deconvoluted peaks (blue) are compared.

### Hydrodynamic behavior of RBD forms analyzed by SEC-HPLC

SEC-HPLC experiments of RBD produced in *P. pastoris* confirmed the absence of aggregated forms and showed a peak compatible with two species between 45-25 kDa (**Figure 6**). Deconvolution of the chromatogram in two components by fitting to two gaussian curve suggested that ∼60% of the signal comes from a higher molecular weight component (>40 kDa), whereas the rest of the signal ∼40% corresponds to a lower molecular weight component (<30 kDa). It is worth mentioning that the exclusion profile corresponding to RBD produced in HEK-293T cells superimposes with the latter, suggesting a more homogeneous glycosylation of the protein.

### Conformational characterization of RBD forms analyzed by *circular dichroism*, fluorescence and thermal-induced unfolding

RBD produced in mammalian and *P. pastoris* cells showed superimposable far-UV circular dichroism (CD) spectra (**Figure 7A**), suggesting a similar secondary structure. Moreover, the CD spectra are identical to that observed for RBD from SARS-CoV-1 produced in yeast^25^. However, given the particular shapes of the *P. pastoris* and HEK-293T SARS-CoV-2 RBD far-UV CD spectra (which show a single minimum at 206 nm and a maximum at 230 nm, the latter suggesting the contribution of aromatic residues to the spectra), it is difficult to estimate the secondary structure content by using standard sets of spectra.

**Figure 7.**
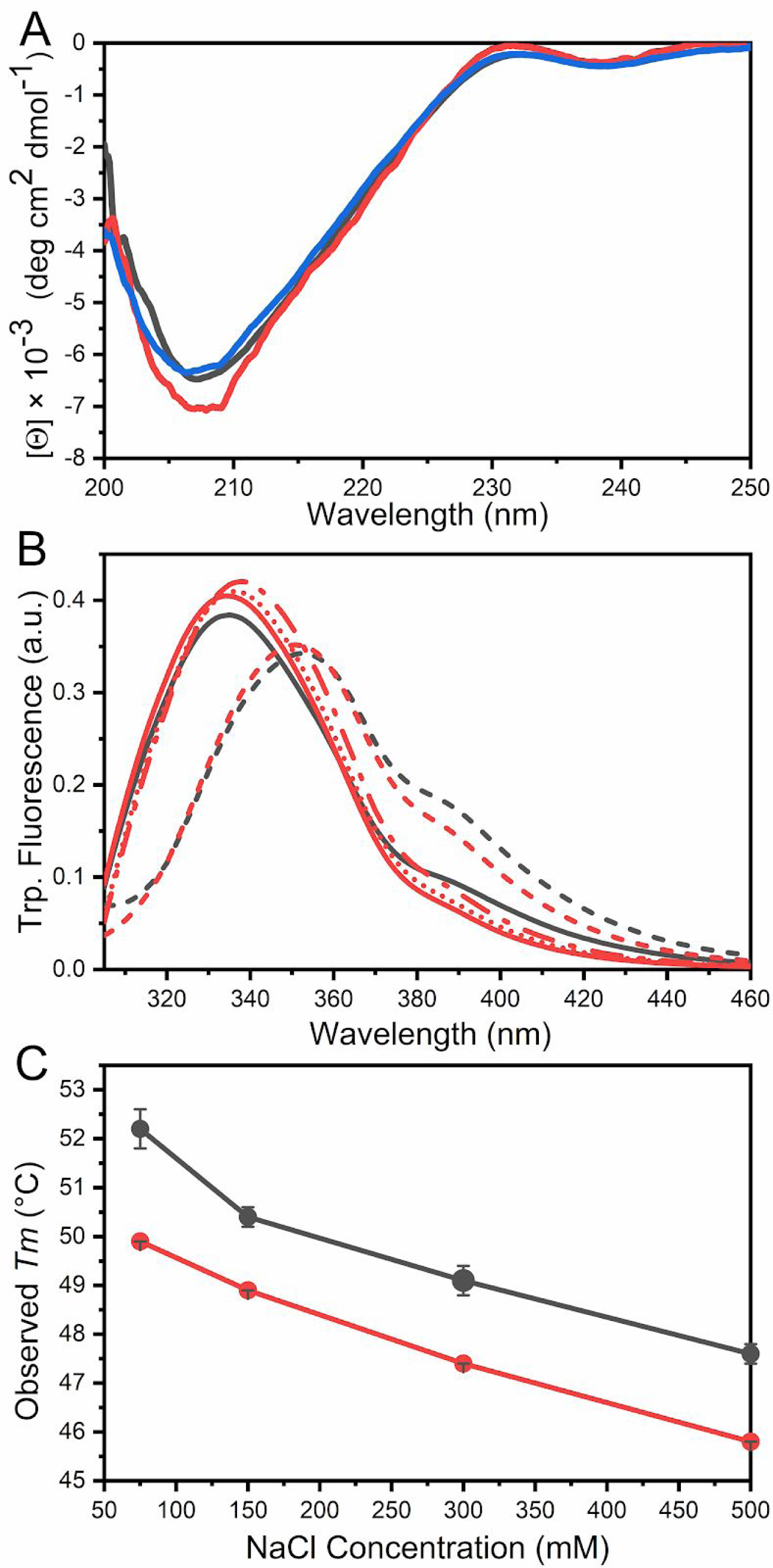
Conformation and stability of Different Purified RBD Forms characterized by Circular Dichroism Spectroscopy, Tryptophan Fluorescence and Temperature-induced Denaturation. (A) Far-UV CD spectra of RBD produced in HEK-293T cells (black), and two different preparations of RBD produced in *P. pastoris* (red and blue). (B) Tryptophan fluorescence emission was monitored by excitation at 295 nm in 20 mM Tris-HCl, 100 mM NaCl, pH7.0 at 25 °C. The spectra of RBD obtained in HEK-293T (black) and in *P. pastoris* (red) are shown, in native conditions (solid line) and in the presence of 4.0 M GdmCl (dashed line) after a 3h incubation. Refolding of RBD produced in *P. pastoris* was performed by dilution to final concentrations of 0.7 M (red dot line) and 1.0 M (red dash-dot line) GdmCl. (C) Stability analysis of RBD. Temperature-induced denaturation of RBD produced in *P. pastoris* (red) and HEK-293T cells (black) under different ionic strength conditions (75, 150, 300 and 500 mM NaCl) was followed by Sypro-orange fluorescence.

We further studied the conformation of RBD produced in HEK-293T cells or *P. pastoris* by tryptophan fluorescence spectroscopy. Spectra corresponding to the native forms of RBD superimposed very well, suggesting that these aromatic residues are located in similar, apolar environments, as inferred by the maximal emission wavelengths observed (337 nm) (**Figure 8B**). The addition of 4.0 M GdmCl resulted in a red shift to 353-354 nm, a result compatible with the exposure of the aromatic side chains to the solvent, and the total unfolding of the protein forms. Interestingly, unfolding of RBD produced in *P. pastoris* showed reversibility when 4.0 M GdmCl was diluted to 0.7 or 1.0 M, as judged by the blue shift from 353 to 337 nm and form 353 to 342 nm, respectively. No reducing agents (*e.g.* DTT, 2-mercaptoethanol) were added to the protein sample, indicating that the dimension of the conformational space corresponding to the unfolded state of RBD was constrained by the native disulfide bonds.

**Figure 8.**
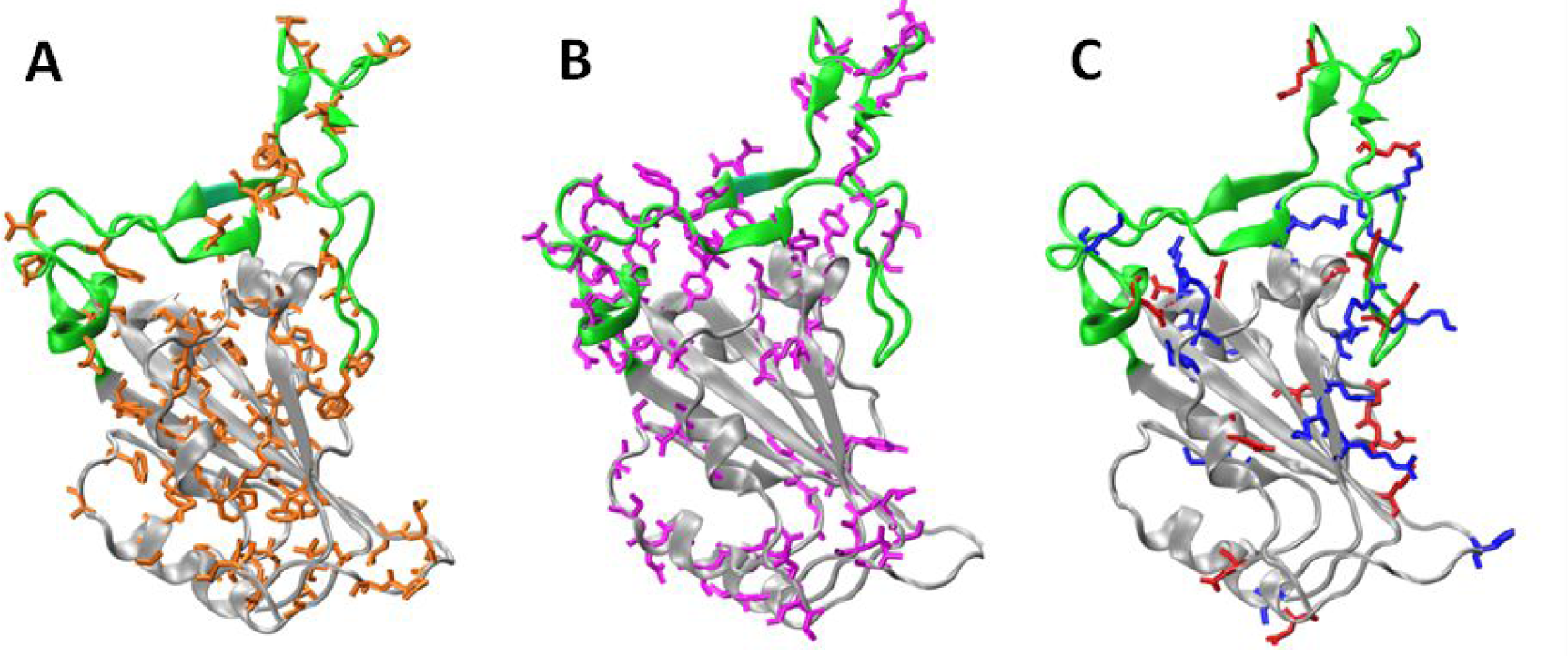
Subdomains and distribution of residue types on RBD. The Core (gray) and the RBM (green) regions are shown. Panels A, B, and C, shows the non-polar residues (orange: A, C, G, I, L, M, F, P, W and V), polar (violet: N, Q, S, T and Y), and charged residues (blue: basic K, R and H, red: acid D and E), respectively. To build the models we used the chain E of pdb structure 6m0j.

The conformational stability of different RBD forms was studied through temperature unfolding experiments. Unfolding was monitored by fluorescence of Sypro-orange, an extrinsic probe that preferentially binds to proteins when they are in unfolded conformations. In these experiments, the observed *T*_m_ usually correlates with *T*_m_ obtained from differential scanning calorimetry experiments. RBD produced in *P. pastoris* consistently showed a slightly lower *T*_m_ value relative to that of RBD from HEK-293T cells (in all assessed conditions), an observation compatible with a reduced resistance to temperature-induced denaturation, which likely reflects a marginally lower conformational stability of RBD produced in *P. pastoris* (**Figure 7C**). When the unfolding process was studied at different ionic strengths, a significant increase in *T*_m_ was observed when NaCl concentration was reduced from 500 to 75 mM (**Figure 7C**), suggesting that the tertiary structure of RBD is stabilized by ionic pair interactions.

### Computational analysis of RBD structure

The dependence of the observed *T*_m_ on the NaCl concentrations, led us to hypothesize that increasing the ionic strength destabilizes RBD conformation by shielding key charged residues. Although the RBD structure suggests some energetic frustration, given that there are several clusters of positively charged residues on RBD surface ((1) R136, R139 and K140; (2) K126, R28, R191; (3) R37, K38, R39 and R148; (4) R85, R90 and K99)^19^, our results suggest that these repulsive interactions are most likely compensated even in the context of the isolated RBD domain (*i.e.* without the rest of Spike or the ACE2 receptor). This would explain why increasing ionic strength has a major destabilizing effect.

The RBD crystallographic structure analysis indicates that residues have a particular distribution according to their type. The core subdomain (residues 333-442 and 504-526 on the Spike protein) is enriched in non-polar residues, whereas the receptor-binding motif (RBM) subdomain (residues 443-503) is enriched in polar ones. On the other hand, the charged residues are preferentially located close to the interface between the Core and RBM subdomains and form an electrostatic network (**Figure 8**).

Among the 14 positively and negatively charged residue pairs interacting at a distance lower than 6 Å, 8 are in the core subdomain, 5 in the RBM, and 1 pair is between the Core and RBM subdomains (**Table 3**). The existence of 6 ionic pair interactions involving at least one occluded charged residue (D398, E406, D442, R454, D467, and R509) is also remarkable.

**Table 3.**
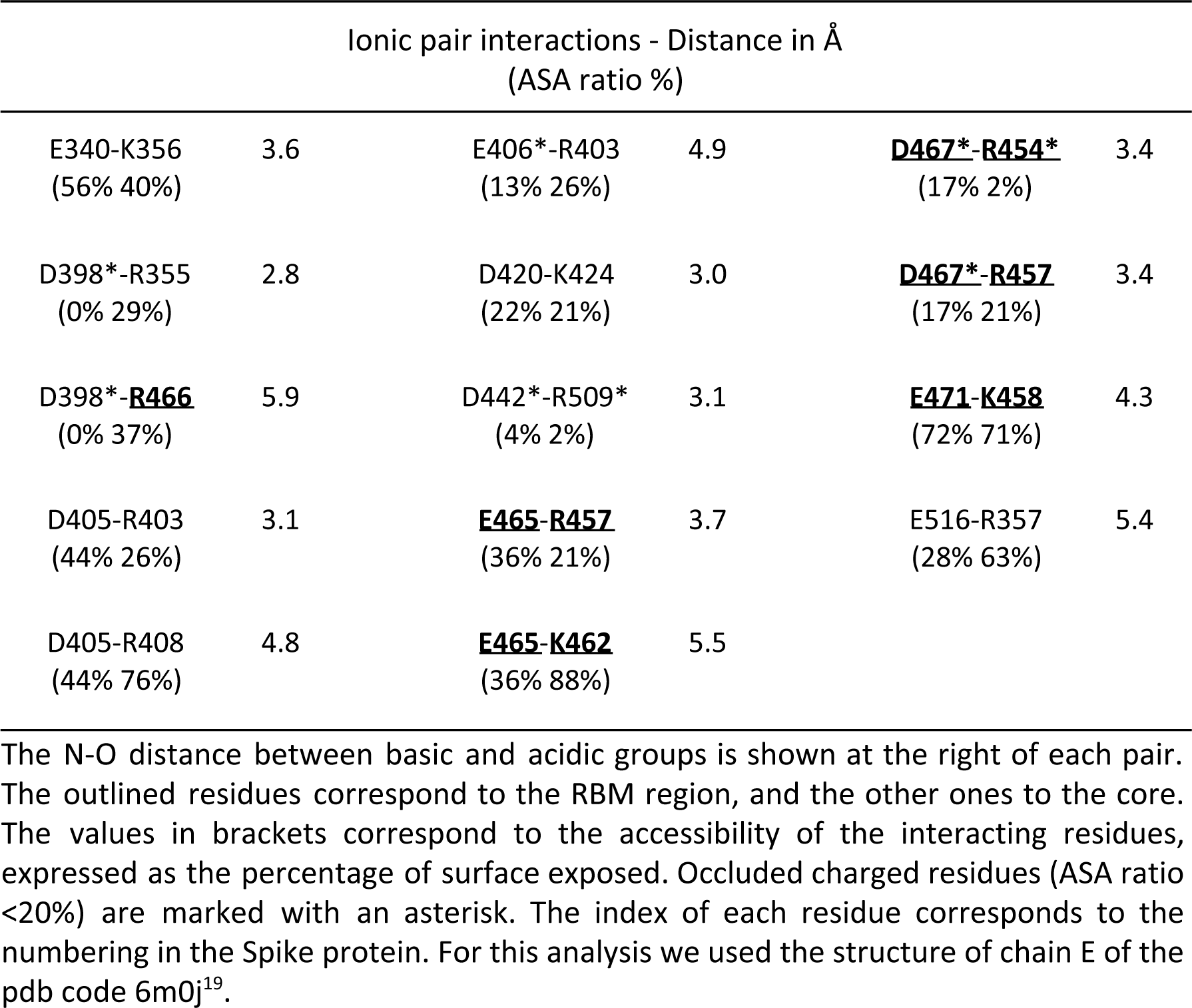
Electrostatic pair interactions between negatively and positively charged residues.

| Ionic pair interactions - Distance in Å<br>(ASA ratio %) |  |  |  |  |  |
| --- | --- | --- | --- | --- | --- |
| E340-K356<br>(56% 40%) | 3.6 | E406*-R403<br>(13% 26%) | 4.9 | **<u>D467\*-R454\*</u>**<br>(17% 2%) | 3.4 |
| D398*-R355<br>(0% 29%) | 2.8 | D420-K424<br>(22% 21%) | 3.0 | **<u>D467\*-R457</u>**<br>(17% 21%) | 3.4 |
| D398*-**<u>R466</u>**<br>(0% 37%) | 5.9 | D442*-R509*<br>(4% 2%) | 3.1 | **<u>E471-K458</u>**<br>(72% 71%) | 4.3 |
| D405-R403<br>(44% 26%) | 3.1 | **<u>E465-R457</u>**<br>(36% 21%) | 3.7 | E516-R357<br>(28% 63%) | 5.4 |
| D405-R408<br>(44% 76%) | 4.8 | **<u>E465-K462</u>**<br>(36% 88%) | 5.5 |  |  |

The N-O distance between basic and acidic groups is shown at the right of each pair. The outlined residues correspond to the RBM region, and the other ones to the core. The values in brackets correspond to the accessibility of the interacting residues, expressed as the percentage of surface exposed. Occluded charged residues (ASA ratio <20%) are marked with an asterisk. The index of each residue corresponds to the numbering in the Spike protein. For this analysis we used the structure of chain E of the pdb code 6m0j<sup>19</sup>.

The importance of these interactions merits further analysis, as they may modulate the conformational dynamics of the RBM, the transitions of the RBD in the Spike trimmer, and/or the interaction with the ACE2 receptor.

### Immune Response elicited by RBD produced in P. pastoris in mice

With the aim of evaluating the ability of the RBD protein produced in *P. pastoris* to stimulate immune response we assessed antibody production in mice by an ELISA assay using plates coated with RBD produced either in HEK-293T or in *P. pastoris*. After a first dose of antigen plus adjuvants, mice presented higher antibody titers than controls, and after a second dose, the levels of antibodies increased significantly in a short period of time (20 days) relative to the first dose (**Figure 9A**). No significant differences in antibody titers were observed between plates coated with RBD from *P. pastoris* or from HEK-293T cells. Thus, immunization of mice with RBD produced in *P. pastoris* plus adjuvant induces a high level of specific IgG class antibodies. Next, Western blots were performed in which RBD produced either in *P. pastoris* or in HEK-293T cells was detected with a serum from mice immunized with RBD produced in HEK-293T cells (kindly provided by Dr. Juan Ugalde, University of San Martín, **Figure 9B, left**), the previously used mouse polyclonal serum from mice immunized with RBD produced in *P. pastoris* (**Figure 9B, middle**), or a primary antibody against the His tag present in both RBD recombinant proteins (**Figure 9B, right**). Similar bands were observed in all three blots.

**Figure 9.**
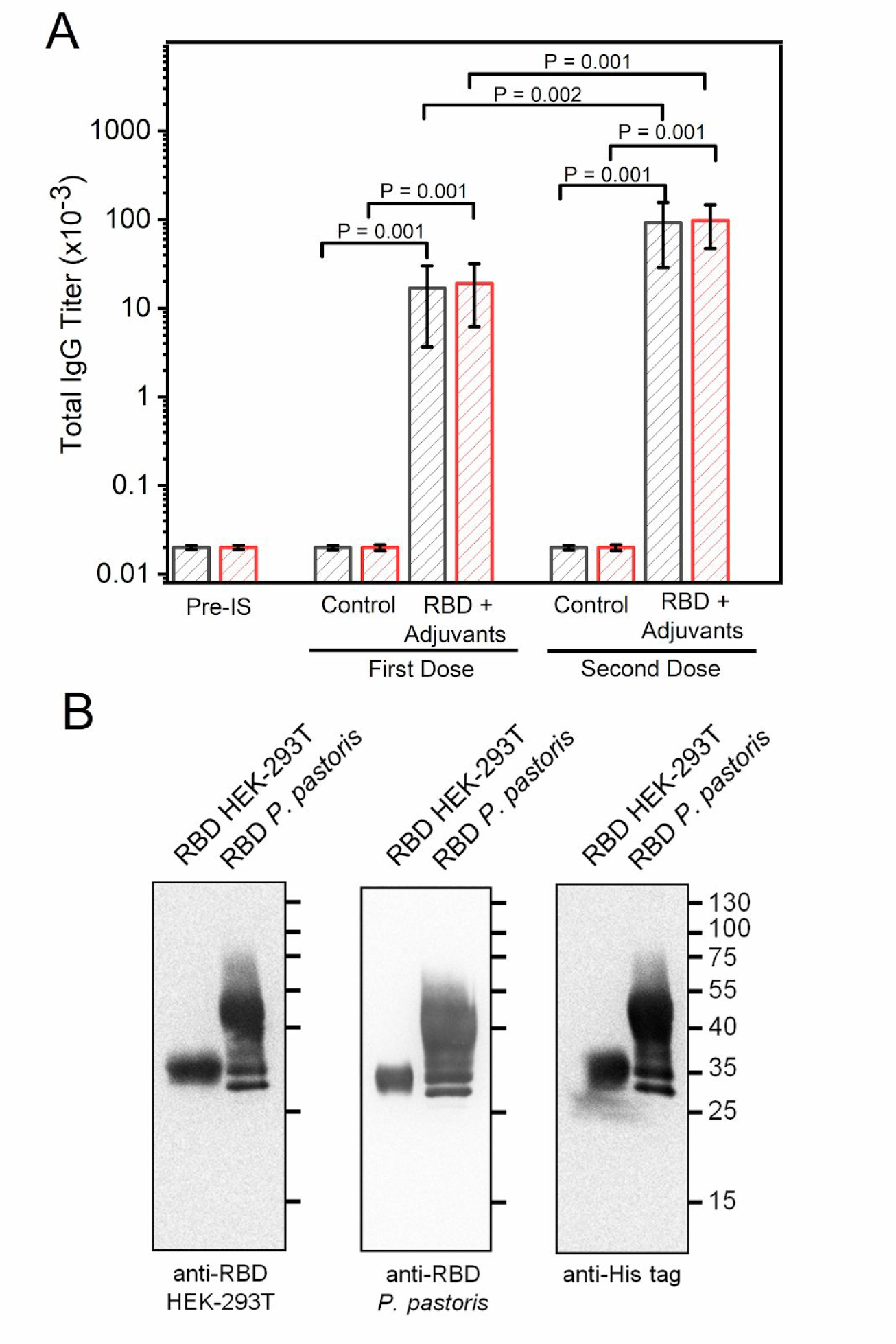
Evaluation of the cross-reactivity of antibodies produced in mice immunized with *P. pastoris* RBD. (A) Titers of antibodies obtained by immunization with RBD from *P. pastoris* plus adjuvants. Each bar represents the group mean (n=5) for specific titers as determined by end-point-dilution ELISA. ELISA was performed with plates coated with RBD protein produced in HEK-293T cells (black sparse bars) or *P. pastoris* (red sparse bars). First dose corresponds to blood samples obtained 30 days post-first immunization, and second dose to samples obtained 20 days post-second immunization. Pre IS, Pre Immune Sera; RBD + Adjuvants, RBD produced in *P. pastoris* + Al(OH)_3_ + CpG-ODN 1826; Control, Al(OH)_3_ + CpG-ODN 1826. P values indicate significant differences between different groups. Bars indicate SD. P values (t-test) are shown for statistically significant differences (p < 0.05). (B) Purified RBD produced in HEK-293T (1.0 *μ*g) and in *P. pastoris* (3.0 *μ*g) were analyzed by Western blot using sera from mice immunized with RBD produced in HEK-293T (anti-RBD HEK-293T, left), or in *P. pastoris* (anti-RBD, *P. pastoris* center). As a control a primary antibody against the His tag present in both RBD recombinant proteins was used (right).

### Production of RBD by fermentation in bioreactor

The fermentation of *P. pastoris* in a 7 L stirred-tank bioreactor for the production of recombinant RBD was carried out using a four-phase procedure described in Methods. In the batch phase, cell concentration reached a maximum level of 15.7 g DCW/L after 18 h of cultivation (**Figure 10**). At this stage, *P. pastoris* exhibited a maximum specific growth rate (*μ*_max_) of 0.21 h and a biomass yield coefficient (*Y*_x/s_) of 0.39 g DCW/g of glycerol. After a spike of dissolved oxygen, the glycerol fed-batch phase was initiated by regulating the feeding in response to the level of dissolved oxygen (DO %). Glycerol feeding was maintained for 22 h, when biomass concentration reached a value of 60.4 g DCW/L. After glycerol feeding was stopped, the transition stage was performed by feeding with a glycerol (600 g/L):methanol (3:1) mixture for 5 h, to allow a slow cell adaptation for the efficient utilization of methanol. At the end of this stage the biomass level reached 63.2 g DCW/L. Next, the methanol fed-batch phase was initiated to induce recombinant RBD expression, through regulation of pure methanol feeding according to DO %. After 48 h of methanol induction and a total fermentation time of 93 h, the culture reached a biomass concentration of 75.3 g DCW/L, which produced a final RBD yield of 45.0 mg/L (>90% pure). Beyond that time point there was no significant change in cell concentration and antigen expression level (data not shown). At the end of the fermentation, a final volume of 5.5 liters of culture was reached, so that the total amount of RBD obtained was 247.5 mg, the volumetric productivity was 0.48 mg/L h, and the total productivity 2.66 mg/h.

**Figure 10.**
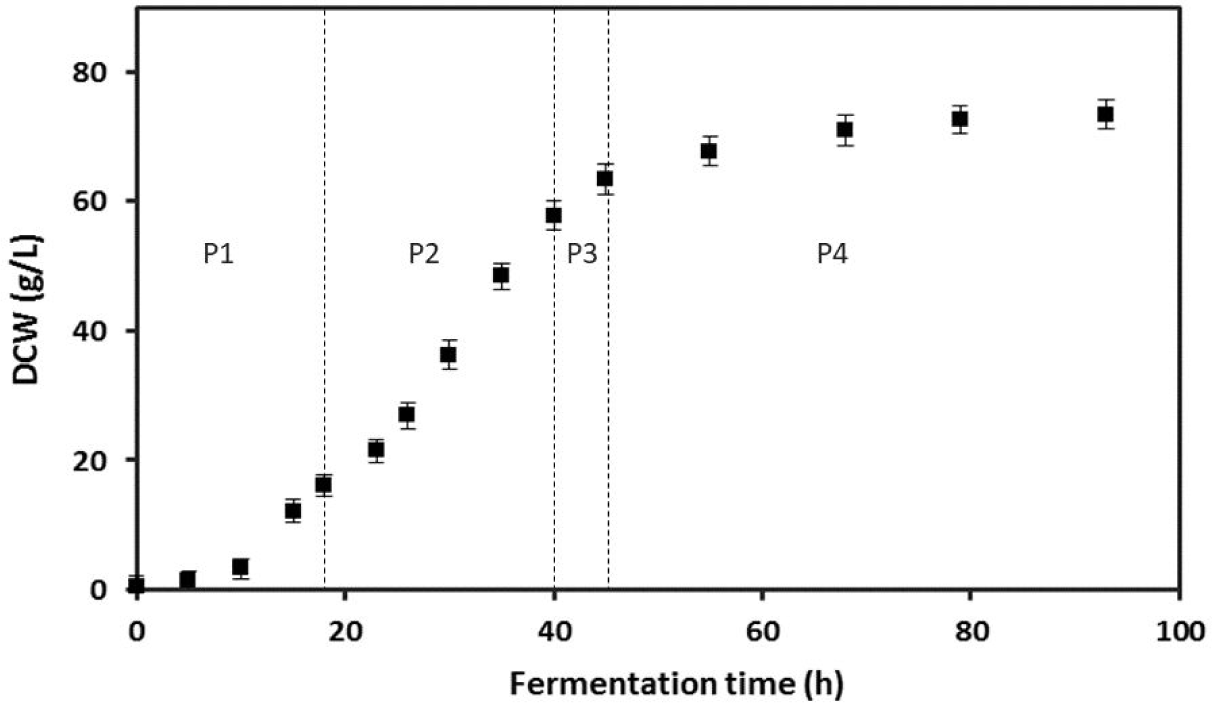
*P. pastoris* biomass concentration (g DCW/L) evolution during bioreactor fermentation. P1: batch phase in LSBM glycerol 40 g/L, P2: Fed-batch phase with 600 g/L glycerol solution, P3: Adaptation phase with glycerol (600 g/L):methanol (3:1) mixture, P4: Induction phase with methanol as the sole carbon source. Error bars indicate 2SD.

## Discussion

This work materialized the first two goals of our consortium assembled to fight COVID-19 pandemia: (a) to express and characterize RBD from SARS-CoV-2, and (b) to produce RBD at low cost with high yield. We were able to express this protein in two different systems: *P. pastoris* and mammalian cells (HEK-293T), which allowed us to gain useful insights concerning RBD conformation and stability.

We attempted to express RBD in *E. coli*, even though an examination of its structure suggested that this system would not be suited for its expression due to the existence of 4 disulfide bonds and a non-globular shape. The *E. coli* SHuffle expression system only yielded insoluble RBD (in inclusion bodies) as expected, which was not further characterized as it was unsuitable for downstream applications (data not shown). In agreement with our results, in a previous attempt to express the similar RBD from SARS-CoV-1 Spike in *E. coli*, this protein was also found in the insoluble fraction, and neither its fusion to thioredoxin, nor to maltose-binding protein, increased its solubility. Moreover, while tagging the protein with glutathione *S*-transferase (GST) increased its solubility^34^, it remained strongly bound to the bacterial chaperone GroEL even after affinity purification^35^, thus making it unsuitable for downstream applications.

By contrast, RBD expression in HEK-293T and *P. pastoris* eukaryotic cells produced soluble and properly folded polypeptides. Their UV-absorption, CD and Trp-fluorescence spectra showed high similarity with those previously described for SARS-CoV-1 RBD produced in *P. pastoris*^25^. RBD expressed in both eukaryotic systems was also characterized by controlled proteolysis and mass spectrometry analysis. Remarkably, the peptide of sequence FPNITNLCPFGEVFNATR was not easily detected by mass spectrometry. A plausible and straightforward explanation for this result is that glycosylation of this peptide at positions N331 (NIT) and N343 (NAT) increases its mass, and thus takes the peptide out of the range of mass analysis. The sequence coverage corresponding to a similar N-terminal peptide of RBD produced in HEK-293T cells was also lower than expected as the peptide VQPTESIVR was not detected, which might be due to the presence of *O*-glycosylations in this stretch.

Proteins translocated to the ER are *N*-glycosylated cotranslationally, and exactly the same glycan Glc_3_Man_9_GlcNAc_2_ is transferred to the N residue in the consensus sequence NXS/T (where X cannot be P) of mammalian, plant and yeast proteins. Glc residues are immediately removed in the ER, where glycans play a key role in the solubility of the glycoproteins and in the so-called “quality control of glycoprotein folding”^36^. Cycles of glucosylation and deglucosylation occur in the endoplasmic reticulum (ER) until the glycoprotein is folded and continues the transit through the secretory pathway or, if unable to fold properly, is retrotranslocated to the cytosol and degraded by the proteasomes. This mechanism guarantees that only properly folded proteins are secreted. Once glycoproteins leave the ER, *N*-glycans are remodeled in the transit through the golgi apparatus and acquire glycan structures that are species specific. In yeast *N*-glycans in mature proteins are of high mannose type while in mammalian proteins are of complex or hybrid type^37^. *O*-glycosylation of S or T residues may also occur in the Golgi, but in this case the monosaccharides are added step by step, and are species specific. When RBD purified from *P. pastoris* and HEK-293T cells were treated with PNGaseF, which removes both high mannose and complex

*N*-glycans, a similar band of ∼26 KDa was observed by SDS-PAGE for both proteins, in agreement with the expected molecular weight of the proteins lacking *N*-glycans. However, only in the case of RBD obtained from *P. pastoris* the same band was observed when proteins were treated with EndoH, which removes high mannose glycans, indicating that RBD from *P. pastoris* bears only the expected high mannose *N*-glycans, while RBD from HEK-293T likely bears complex or hybrid ones. It has been previously reported that SARS-CoV-2 RBD is produced as two predominantly *N*-glycosylated forms of∼34 and ∼27 kDa when expressed in Sf9 insect cells^23^.

Exhaustive removal of *N*-glycans RBD obtained from HEK-293T cells produced two protein forms that migrated as distinct bands, suggesting the presence of either two *O*-glycosylated isoforms, or of two peptide isoforms. Shajahan and coworkers have evaluated the *O*-glycosylation of SARS-CoV-2 Spike protein produced in HEK-293 cells by searching LC-MS/MS data for common *O*-glycosylation modifications. These authors found *O*-glycosylation sites at positions T323 and S325, which provides further support to our suggestion^37^.

For secretory recombinant proteins produced in yeast, high mannose hyperglycosylation may be a major issue, as it may potentially alter functional properties of the proteins^26^. However, even though RBD expressed in HEK-293T and *P. pastoris* exhibited different glycosylation patterns, both conformational and stability studies carried out in the present work suggest that both polypeptides have similar structures. The stability of both forms was similarly sensitive to changes in ionic strength, a result in good agreement with our computational modelling, which predicts that a set of ionic interactions stabilizes RBD structure and most likely modulates its internal motions. Interestingly, when RBD was unfolded in the absence of reducing agents the process was reversible, as judged by the analysis of Trp fluorescence spectra, which suggests that Trp residue emission occurs in an apolar and likely more rigid environment upon refolding.

Previously, the thermal stability of a mutated version of RBD from SARS-CoV-1 Spike named RBD219-N1 (residues 319-536 from Spike, where the first Asn of RBD, residue 318, was deleted to avoid glycosylation at the N-terminal region of the protein) was analyzed by thermal shift monitoring of extrinsic fluorescence^38^. Remarkably, the RBD219-N1 denaturation profile showed an average melting temperature of approximately 57 °C, a value significantly higher than that observed for RBD from SARS-CoV-2 in this work (approximately 50 °C), which might be due to the fact that denaturation of RBD219-N1 was carried out at a considerably lower ionic strength^38^. In addition, the pH of the protein sample was not constant throughout the experiment, given that the p*K*a of the Tris buffer used in this work is highly temperature dependent.

Alternatively, the suppression of the N-terminal glycosylation in SARS-CoV-1 RBD might have an effect on its conformational stability. Nevertheless, the possibility that SARS-CoV-1 RBD has particular features that might increase its stability relative to that of SARS-CoV-2 RBD should not be excluded.

In our hands, RBD in its native state was stable under a broad range of pH and concentrations. Although it exhibited a low tendency to aggregate at high concentrations, no significant complications were observed during filter-protein concentration or dialysis, which was performed either to change the buffer or to remove imidazole after protein purification. Freezing (-80 °C) and subsequent thawing of RBD did not result in protein aggregation at protein concentrations of 30-40 *μ*M or lower, therefore this strategy was used for its storage, as it made unnecessary the use of stabilizing molecules such as glycerol or trehalose. However, it should be noted that RBD precipitation was occasionally observed after thawing, at protein concentrations above 80 *μ*M.

*P. pastoris*-produced RBD was able to stimulate antibody production in mice, and the resulting immune sera were capable of detecting RBD produced not only in *P. pastoris* but also in HEK-293T cells. Finally, scaling up of RBD expression in *P. pastoris* could be performed in a bioreactor with yields greater than 45 mg L^-1^, which potentially allows the large scale immunization of animals in order to produce neutralizing antibodies, or the development of SARS-CoV-2 vaccines. Future biotechnological developments will be facilitated by the inclusion of a Sortase-A enzyme recognition site within the RBD coding sequence, which allows the native covalent coupling of RBD to fluorescent probes, peptides, proteins, or modified surfaces, through an efficient transpeptidation reaction^39^. Thus, the Sortase-A-mediated transpeptidation will allow future efficient *in vitro* covalent linking of RBD with protein carriers independently-produced in low cost systems such as *E. coli*.

## Methods

### Expression of RBD in mammalian cells

For RBD expression in mammalian cells, a DNA fragment optimized for expression in human cells encoding RBD (Spike residues from 319 to 537, preceded by the IL-2 export sequence (MYRMQLLSCIALSLALVTNS) and followed by a C-terminal Sortase-A recognition sequence for covalent coupling^39^ and a His6 tag for purification (LPETGHHHHHH) was synthesized by GenScript (NJ, USA) and cloned into the pCDNA3.1(+) plasmid vector (ampicillin R). Expression of RBD was carried out in the HEK-293T cell line kindly provided by Xavier Saelens (VIB-University of Ghent, Belgium). HEK-293T cells were grown in high glucose (4.5 g L^-1^ glucose) Dulbecco’s modified Eagle’s medium (DMEM, Thermo Fisher Scientific) supplemented with 10% fetal bovine serum (FBS, Natocor), penicillin/streptomycin (100 units mL^-1^ and 100 µg mL^-1^ respectively, Thermo Fisher Scientific) and 110 mg L^-1^ of sodium pyruvate (Thermo Fisher Scientific) in a 37°C humidified incubator containing 5% CO_2_. Cells were plated (2 x 10^7^ cells per 150 mm plate) and grown for 24 h before transfection with Polyethylenimine (PEI, Sigma) according to the manufacturer’s instructions. Cells were grown for 72h before harvesting the culture medium.

Mammalian cell culture medium was centrifuged twice at 12,0000xg for 20 min at 4 °C, later the supernatant pH was adjusted to 8.0 with equilibration buffer (50 mM sodium phosphate, 300 mM NaCl and 20 mM imidazole, pH 8,0). RBD was purified using a previously equilibrated Ni^2+^-NTA-agarose column. RBD was eluted by increasing concentrations of imidazole prepared in the equilibration buffer. Fractions containing the recombinant protein, as judged by the SDS-PAGE analysis, were pooled and dialyzed against a buffer without imidazole (20 mM Sodium phosphate, 150 mM NaCl, pH= 7,4).

### Expression of RBD in Pichia pastoris

The RBD coding sequence with codon optimization for *P. pastoris* (Spike amino acid residues 319-537) fused to the *Saccharomyces cerevisiae* alpha factor secretion signal (N-terminal)^40^ followed by a C-terminal Sortase-A recognition sequence and a His6 tag (C-terminal) was synthesized and cloned into pPICZalpha by GenScript (NJ, USA) using EcoRI and SacII restriction sites to produce pPICZalphaA-RBD-Hisx6.

The SacI linearized pPICZalphaA-RBD-Hisx6 vector (10 *μ*g) was used to transform electrocompetent X-33 *P. pastoris* strain at 2.5 kV, 25 uF, 200 ohm. Cells recovered in ice-cold 1.0 M sorbitol (Sigma) were plated in YPDS (YPD: 1% yeast extract, 2% bactopeptone (Difco), 2% glucose, plus 1.0 M sorbitol) supplemented with 100 μg mL^-1^ zeocin (Invitrogen) and incubated four days at 28 °C. Selected colonies (45) were transferred to increasing amounts of zeocin (100 to 500 μg mL^-1^). The integration at AOX site in clones that were resistant to the highest amount of zeocin was confirmed by colony PCR using primers AOX_for_ (5’-GACTGGTTCCAATTGACAAGC-3’) and RBD_rev_ (5’ GTTCCATGCAATGACGCATC 3’).

For RBD production in batch, single colonies were used to inoculate BMGY medium (1% yeast extract, 2% bactopeptone, 1.34% YNB, 400 μgL^-1^ biotin, 0.1 M potassium phosphate, pH 6.0, and 1% glycerol) and cultures were grown at 28 °C with agitation at 250 rpm until the culture reached a OD_600nm_= 4-6. Cells were harvested, resuspended in either Buffered Methanol-complex Medium (BMMY: 1% yeast extract,

(Difco) 2% peptone, 100 mM potassium phosphate pH 6.0, 1.34% YNB, 0.4 mg L^-1^ biotin, 0.5 % methanol, Sintorgan) or buffered minimal methanol medium containing histidine (Sigma) (BMMH: 100 mM potassium phosphate pH 6.0, 1.34% YNB, 0.4 mg L^-1^ biotin, 4 mg L^-1^ histidine, 0.5 % methanol) to an initial DO_600nm_= 1.0, and incubated at 28 °C with shaking at 250 rpm in flasks covered with microporous tape sheets for better oxygenation. Every 24 hs methanol was added to a final concentration of 0.5% and pH was adjusted to 6 if necessary. Induction was maintained during 72-90 h at 28 °C, then cells were removed by centrifugation at 3000 ×g for 10 min and the supernatant was frozen at -80C until it was used.

The purification of RBD from culture media was performed using a NTA-Ni^2+^ column previously equilibrated with 50 mM Tris-HCl, 150 mM NaCl, 10% glycerol, pH 8.0 (equilibration solution). The media supernatants was adjusted to pH 8.0 with NaOH, centrifuged 20 min at 12,000 xg and loaded to the column. The flow through was reloaded twice. The column was washed with an equilibration solution containing 20-30 mM imidazole. Finally, RBD was eluted with an equilibration solution containing 300 mM imidazole. The purified protein was dialyzed twice in 20 mM Tris-HCl, 150 mM NaCl buffer, pH 7.4, and quantified by absorbance at 280 nm (see below) and stored at -80 °C.

### UV Absorption Spectroscopy

The concentration of recombinant RBD was determined by UV spectrophotometry, using the following extinction coefficients derived from the protein sequence (considering all the disulfide bonds formed): for RBD produced in *P. pastoris*: ε_280nm_= 33850 M^−1^ cm^−1^ (Abs _280nm_= 1.304 for a 1 mg mL^-1^ protein solution); for RBD produced in HEK-293T cells (without considering IL2 export signal sequence: ε_280nm_= 33850 M cm (Abs 280= 1.300 for a 1 mg mL^-1^ protein solution).

Absorption spectra (240-340 nm range, using a 0.1-nm sampling interval) were acquired at 20 °C with a JASCO V730 BIO spectrophotometer (Japan). Ten spectra for each sample were averaged, and blank spectra (averaged) subtracted. A smoothing routine was applied to the data by using a Savitzky-Golay filter and subsequently the 4th derivative spectra were calculated.

### SDS-PAGE and Western Blotting Analysis

Purified RBD produced in *P. pastoris* or in HEK-293T cells was boiled in sample buffer (4% SDS, 20% glycerol, 120 mM Tris, pH 6.8, 0.002% bromophenol blue, 200 mM 2-mercaptoethanol) and separated in 12% SDS-PAGE. Proteins were either stained with Coomassie brilliant blue G-250 or transferred to nitrocellulose membranes (GE Healthcare). The membranes were blocked with 5% milk in 0.05% Tween TBS at room temperature for 1 h and then incubated at 4 °C overnight with a specific polyclonal serum produced by immunization of mice with RBD produced in HEK-293T cells. HRP-conjugated anti-mouse were incubated for 1 h at room temperature and visualized by enhanced chemiluminescence (ECL, Thermo Scientific).

### Circular Dichroism and Tryptophan Fluorescence

CD spectra measurements were carried out at 20 °C with a Jasco J-815 spectropolarimeter. Far-UV and near-UV CD spectra were collected using cells with path lengths of 0.1 and 1.0 cm, respectively. Data was acquired at a scan speed of 20 nm min^-1^ (five scans were averaged, scans corresponding to buffer solution were averaged and subtracted from the spectra). Values of ellipticity were converted to molar ellipticity.

Steady-state tryptophan fluorescence measurements were performed in an Aminco-Bowman Series 2 spectrofluorometer equipped with a thermostated cell holder connected to a circulating water bath set at 20 °C. A 0.3-cm path length cell was used. The excitation wavelength was set to 295 nm and emission data were collected in the range 310–450 nm. The spectral slit-width was set to 3 nm for both monochromators. The protein concentration was 3.0-5.0 μM, and the measurements were performed in a buffer containing 20 mM sodium phosphate, 150 mM NaCl, pH 7.40.

### Temperature-Induced Denaturation Monitored by Sypro-Orange

Temperature-induced denaturation of RBD forms was monitored by the change in the Sypro Orange dye (Thermo Fisher) fluorescence using protein at a 5.0 μM concentration in 50 mM sodium phosphate buffer, pH 7.0. Samples without protein were also included as controls. The dye was used at 2 × (as suggested by Thermo Fisher Scientific). The temperature slope was 1 °C min^-1^ (from 20 to 90 °C). Excitation and emission ranges were 470–500 and 540–700 nm respectively.

The fluorescence signal was quenched in the aqueous environment but became unquenched when the probe bound to the apolar residues upon unfolding. Experiments by triplicate were carried out in a Step One Real-Time-PCR instrument (Applied Biosystems, CA, U.S.A.).

### Hydrodynamic Behavior by SEC-HPLC

SEC-HPLC was performed using a Superose-6 column (GE Healthcare). The protein concentration was 20-30 μM, a volume of 50μL was typically injected, and the running buffer was 20 mM Tris-HCl, 100 mM NaCl, 1 mM EDTA, pH 7.0. The experiment was carried out at room temperature (∼25 °C) at a 0.5 mL min^-1^ flow rate. A JASCO HPLC instrument was used. It was equipped with an automatic injector, a quaternary pump and a UV-VIS UV-2075 (elution was monitored at 280 nm).

### Production of RBD in P. pastoris by bioreactor fermentation

*P. pastoris* was grown on a solid YPD medium containing 20 g L^-1^ peptone, 10 g L^-1^ yeast extract, 20 g L^-1^ glucose and 20 g L^-1^ agar medium at 30 ± 1 °C. As previously described by Chen and coworkers^38^, liquid cultivation was carried out using low salt medium (LSM) containing 4.55 g L^-1^ potassium sulfate, 3.73 g L^-1^ magnesium sulfate heptahydrate, 1.03 g L^-1^ potassium hydroxide, 0.23 g L^-1^ calcium sulfate anhydrous, 10.9 mL L^-1^ phosphoric acid 85% and 40 g L^-1^ glycerol, in order to prevent salt precipitation during downstream processing due to increased pH. After sterilization, 3.5 mL per liter of filtered biotin solution (0.02 % w/v) and 3.5 mL per liter of trace metal solution (PTM1) were added. PTM1 contained per liter: 6.0 g copper (II) sulfate pentahydrate, 0.08 g sodium iodide, 3.0 g manganese sulfate-monohydrate, 0.2 g sodium molybdate-dihydrate, 0.02 g boric acid, 0.5 g cobalt chloride, 20.0 g zinc chloride, 65.0 g ferrous sulfate-heptahydrate, 0.2 g biotin and 5 mL sulfuric acid 5.0 mL.

Fermentations were conducted in a stirred-tank bioreactor using a four-stage process based on Celik *et al*.^26, 41^, with slight modifications. The first stage consisted in a batch culture using LSM medium with 40 g L^-1^ glycerol as carbon source and supplemented with 3.5 mL L^-1^ PTM1 and 3.5 mL L^-1^ biotin solution (0.02 % w/v). Under these conditions yeast cells grew and reached high biomass levels without expression of RBD, as it was under the control of *AOX1* promoter, which is repressed when glycerol is provided as an unlimited substrate. In the second phase, a glycerol solution (600 g L^-1^ solution supplemented with 12 mL L^-1^ PTM1) was fed into the culture at a growth-limiting rate to increase cell concentration. At the same time, a gradual derepression of *AOX1* promoter takes place under this glycerol-limited condition. This phase was initiated after glycerol depletion was evidenced by an oxygen spike. Glycerol feeding was automatically regulated according to the percentage of dissolved oxygen (DO %) in the culture, with a cut-off of 60 % saturation. Subsequently, a short transition stage was conducted by feeding a glycerol:methanol (3:1) mixture, thus allowing the adaptation of cells for the growth in the presence of methanol. Finally, the induction stage was carried out by adding pure methanol (supplemented with 12 ml L^-1^ PTM1) in a fed-batch mode with a growth-limiting rate. Methanol feeding was also automatically regulated according to the level of DO % in the culture, with a cut-off 50 % saturation.

In order to obtain the inoculum for bioreactor fermentations, transformed *P. pastoris* cells grown on YPD agar plates were inoculated into a 100-mL flask containing 20 mL of LSM medium with 10 g L^-1^ glycerol (supplemented with PTM1 and biotin) and cultured overnight at 30 ± 1°C. A volume of 300 mL of LSM containing 10 g L^-1^ glycerol (supplemented with PTM1 and biotin) in a 1 L Erlenmeyer flask was inoculated with the overnight culture and incubated at 30 ± 1°C until the culture reached an OD_600_ of ∼16. This culture was used to inoculate 3 L of LSM with 40 g L^-1^ glycerol (supplemented with 3.5 mL L^-1^ PTM1 and 3.5 ml L^-1^ biotin 0.02% w/v) in a 7-L BioFlo 115 bioreactor (New Brunswick Scientific; Edison, NJ), which was interfaced with Biocommand Bioprocessing software (New Brunswick Scientific) for parameter control and data acquisition.

Temperature was maintained at 30 ± 1°C throughout batch and glycerol fed-batch phases and at 25 ± 1°C during transition and induction stages. The pH was maintained at 5.0 during the first two phases, and at 5.5 in the last two, by adding H_3_PO_4_ (42.5%) and 14% (v/v) NH_4_OH, which also served as a nitrogen source. DO % was regulated by an agitation cascade (maximum of 1200 rpm) and supplemented with filter-sterilized (0.22 μm) air and pure oxygen when needed. The pH was measured using a pH electrode (Mettler-Toledo GmbH, Germany), and the oxygen concentration was measured with a polarographic probe (InPro6110/320, Mettler-Toledo GmbH). Foam formation was avoided by the addition of 3 % (v/v) antifoam 289 (Sigma-Aldrich; St. Louis, MO). Samples were withdrawn throughout the fermentation process with the purpose of evaluating the biomass and recombinant protein expression.

The optical density of *P. pastoris* culture samples was measured at 600 nm using an UV-Vis spectrophotometer and converted to dry cell weights (DCW, in g L^-1^) with a previously calculated DWC versus OD_600nm_ calibration curve in accordance with the formula: DCW= 0.269 x OD_600nm_, R = 0.99.

The protein profile throughout the methanol-induction phase was analyzed by 12% SDS-PAGE, and gels were stained with Coomassie brilliant blue G-250. RBD expression was confirmed by Western blot analysis using anti-RBD and anti-his antibodies. Total protein content was estimated by measuring the absorbance at 280 nm using a Beckman spectrophotometer.

### Glycan removal from RBD

RBD (5 μg) produced in HEK-293T cells or *P. pastoris* were denatured 10 min at 100 °C with 0.5% SDS and 40 mM DTT. Then, 1% Nonidet P-40, 50 mM buffer sodium phosphate buffer pH 7.5, and 50 mU of PNGaseF (New England Biolabs) were added to remove complex glycans from RBD produced in HEK-293T cells. High mannose glycans of RBD produced in *P. pastoris* were removed by incubation with 5 mU of EndoglycosidaseH (Roche) in 50 mM sodium citrate buffer, pH 5.5. Reactions were incubated during 1 h at 37 °C and analyzed by SDS-PAGE 12%. Parallel control reactions were performed under the same conditions but without adding the endoglycosidase in each case.

### Bioinformatic Studies

A total of 75355 amino acid sequences from Spike protein were downloaded from the Global Initiative for Sharing All Influenza Data (GISAID) database (https://www.gisaid.org)27. Multiple sequence alignment was obtained using MAFFT v.7453^42^. The RBD region was extracted with the EMBOSS package^43^ using the RBD region of Uniprot accession QHN73795. Protein identity analysis was performed by BLAST using the RBD protein sequence as a query and an e-value of 0.001 was used. Sequence identity and coverage percentages were registered and counted.

### Reverse phase HPLC

For HPLC, a JASCO system equipped with an autoinjector, an oven (thermostatized at 25 °C) and a UV detector was used. A gradient from 0 to 100% acetonitrile was performed (0.05% TFA (v/v) was added to the solvents). An analytical C18 column was used (Higgins Analytical, Inc. U.S.A.), with a 1.0 mL min-1 flow.

### MALDI TOF for intact mass analysis

The protein samples were analyzed using a MALDI TOF TOF mass spectrometer (Applied Biosystems 4800 Plus) operating in linear mode. Previously, the samples were desalted on ZipTip C_4_ column (Millipore, Merck KGaA, Darmstadt, Germany), then eluted in a matrix solution of sinapinic acid 10 mg ml^-1^ in 70 % acetonitrile, 0.1 % TFA or 2,5 dihydroxy benzoic acid 5 mg ml^-1^ in 70 % acetonitrile, 0.1 % TFA and deposited on the MALDI plate. The spots were allowed to dry and finally the samples were ablated using a pulsed Nd:YAG laser (355 nm). Spectra were acquired in positive or negative mode, depending on the sample characteristics.

### Tryptic digestion

The protein samples were digested with trypsin (Promega, mass spectrometry grade) in (NH_4_)HCO_3_ buffer (0.1 M, pH 8.0), after ON at 37 C, the cysteine side chains were previously modified with DTT-iodoacetamide. The tryptic mixture was desalted using a μZipTip μC_18_ column (Millipore, Merck KGaA, Darmstadt, Germany) and were eluted with a saturated matrix solution of α cyano-4-hydroxycinnamic acid in acetonitrile:water (70:30, 0.1% TFA). Alternatively, the samples were digested in a gel following a similar protocol to that used for in solution digestion, with a previous washing step.

### MALDI TOF TOF for tryptic peptides analysis

The spectra were first acquired in reflectron mode and the main signals studied in MS/MS mode. The resulting MS/MS spectra were analyzed using the MASCOT search engine^44^ (Matrix Science) program and COMET^45^ at Transproteomic Pipeline. Also, for the manual analysis of spectra in reflectron mode, the GPMAW (Lighthouse data) program was used.

### Molecular Modelling

The molecular Modelling analysis of the RBD domain was done using the chain E of the pdb structure 6M0J^19^. Figures of this structure were done using VMD^46^. The identification of residues making moderate and strong electrostatic interactions within RBD was performed using the Salt Bridges plug in of VMD. For the analysis we used a 6.0 Å cut-off distance between side chain oxygen and nitrogen atoms of residues D, E, K and R. The accessible surface area calculations for the residues of RBD was done using the GetArea server http://curie.utmb.edu/getarea.html using a 1.4 Å probe radius.

### Immunization protocols

Immunization of mice was carried out by experts from the High Level Technological Service CONICET (STAN No. 4482), under ISO9001 guidelines and those from the Institutional Committee for the Care and Use of Laboratory Animals (CICUAL). BALB/c mice were obtained from the animal facility of the Faculty of Veterinary Sciences, University of La Plata (Argentina), and housed at the animal facility of the Instituto de Ciencia y Tecnología Dr. César Milstein, Fundación Pablo Cassará. Female mice (6-8 week-old) were immunized intraperitoneally with 40 µg RBD protein produced in *P. pastoris* in the presence of the HPLC-grade phosphorothioate oligonucleotide CpG-ODN 1826 (5’ TCCATGACGTTCCTGACGTT 3’) (20 µg/mouse/dose) (Oligos etc. Inc., Integrated DNA Technologies, OR, USA) and aluminum hydroxide (Al(OH)_3_) (20% (v/v)/mouse/dose) and boosted on day 30 with the same dose. Additional control animals were injected with Al(OH)_3_ (20% (v/v)) plus CpG-ODN 1826 (20 µg) per mouse with the same immunization schedule. Pre-immune sera also were collected before starting the immunization. Blood samples were obtained at 30 days post-first immunization (antigen prime) and 20 days post-second immunization (antigen boost) by venipuncture from the facial vein. After coagulation at room temperature for 1-2 h, blood samples were spun in a centrifuge at 3000 rpm/min for 10 min at 4 °C. The upper serum layer was collected and stored at -20°C.

### Identification of serum antibody against protein RBD in mice using an ELISA assay

Standard ELISA procedures were followed to measure antibody response against RBD. Briefly, RBD protein produced in *P. pastoris* or HEK-293T cells was used to coat flat-bottom 96-well plates (Thermo Scientific NUNC-MaxiSorp) at a final concentration of 1 µg/ml (100 µl/well) in phosphate-buffered saline (PBS) coating buffer (pH 7.4) at 4 °C overnight. After blocking with 8% non-fat dry milk PBS for 2h at 37°C the plates were washed 5 times with PBS containing 0.05% Tween 20 (PBST). Serially diluted mouse sera were incubated at 37 °C for 1.5 h in PBS containing 1% non-fat dry milk (blocking solution), and then the plates were washed with PBST. For total specific IgG determination, IgG horseradish peroxidase (HRP)-conjugated antibody (DAKO P0447) was diluted 1/1000 in blocking solution and added to the wells. After incubation for 1 h at 37 °C, plates were washed 5 times with PBST and developed with 3,3’,5,5’-tetramethylbiphenyldiamine (TMB) for 15 min. The reaction was stopped with 50 µl/well of 1.0 M H_2_SO_4_ (stop solution). The absorbance was measured in a microplate reader (Thermo Multiscan FC ELISA) at 450 nm (A_450_). The antibody titer was determined as the inverse of the last dilution that was considered positive, with a cut-off value defined as A_450_= 0.20, which was twice as high as that from a pool of Normal mice Sera (from 30 unimmunized animals). Statistical significance was evaluated by the Student’s t-test, using a logarithmic transformation of the ELISA titers. Differences were considered significant if p < 0.05.

## Acknowledgments

We thank LANAIS-PRO-EM for the support with mass spectrometry analysis of proteins and peptides, and Fundación Ciencias Exactas y Naturales from Universidad de Buenos Aires for their help. We thank Dr. Juan Ugalde from UNSAM for providing a serum from mice immunized with RBD produced in HEK-293T. We would like to specially thank Dr. Diego U. Ferreiro for his initial suggestions concerning SARS-CoV-2 protein expression.

## Author contributions

(names must be given as initials) All authors contributed equally to this work.

## Additional Information

(including a Competing Interests Statement) The author(s) declare no competing interests.

## UniProt Accession IDs

UniProtKB - P0DTC2 (SPIKE_SARS2); Spike glycoprotein from SARS-CoV-2.

